## Supplementary Information for "Minimal exposures reveal visual processing priorities"

### Supplementary Note 1: LCD tachistoscope

#### LCD tachistoscope

We used a custom-made LCD tachistoscope (**Supplementary Fig. 1**), an adapted version of the design described by Sperdin et al.[1,2]. In both versions, two LCD screens are employed with one placed upright vertically and the other placed horizontally, aligned to the top of the other screen. Both screens are branded Philips 223V5LHSB2 with a resolution of 1920 × 1080 pixels, with a size of 476.6 mm in width and 268.11 mm in height, giving a pixel pitch of 0.248 × 0.248 mm. Both screens are fixed in a rigid aluminium frame, which is covered both internally and externally by matte-finished Plexiglass, which in addition to providing structural support also prevents light reflection. Between the two screens, a diagonally placed semi-permeable mirror allows light to pass from the vertical screen while it reflects light emitted from the horizontal screen placed above it. Therefore, both screens are presented to the observer superimposed on each other if their backlights are simultaneously turned on, as when a target stimulus is presented with sub-millisecond precision. This semi-permeable mirror is a Pilkington MirroView™ 50/50 glass of 418 × 504 mm, 6 mm thickness, toughened for robustness (to avoid image deformation due to gravity-caused deflection). We can control which screen is visible to the observer at each time point with a precision of 2±1 microseconds (μs) by controlling the screens' backlights, which are powered by an independent power supply (36V, 108 Watt) with 20 Ohms rheostat in series to dim screen luminosity. The semi-permeable mirror is not perfectly 50:50, resulting in one screen being more luminous than the other. Rheostats are therefore employed in order to decrease the voltage of the more luminous screen, so it matches the screen with lower luminosity and corrects for screen disparities. Unlike the design of Sperdin et al.[1], in our design backlights are controlled by a dedicated micro-controller (ATmega328 AVR) instead of using a parallel port. The micro-controller receives the instruction via serial communication (USB 2.0) to switch on the backlight for the stimulus' predetermined presentation duration. Using a dedicated micro-controller offers two advantages: it prevents hardware compatibility issues since parallel ports are becoming very uncommon; and it provides higher precision and consistency given that its only function is controlling backlight thus it is not affected by computer workload. A minor disadvantage, however, is that the absolute stimulus presentation time is affected by the time it takes for the computer to send the instructions, which is not constant and can take a few milliseconds. Nevertheless, while the onset of stimulus presentation may vary slightly, the exact stimulus presentation duration is controlled with a precision of 2±1 µs for presentations under 16 ms and of 20 µs for stimulus presentations between 16 ms and 10 seconds. The tachistoscope included a signal output to synchronise stimulus presentation durations with other hardware (e.g., EEG signal triggers). In all our experiments, the vertical screen presented all images that did not involve extremely brief exposure durations, including fixation, placeholders, response cue, etc., and its backlight was therefore always on, whereas the horizontal screen presented the stimuli that involved extremely brief exposure durations (described below), and its backlight was therefore off, except during stimulus presentations when it came on for a very brief period; during this period, the content of both screens was visible to the observer.

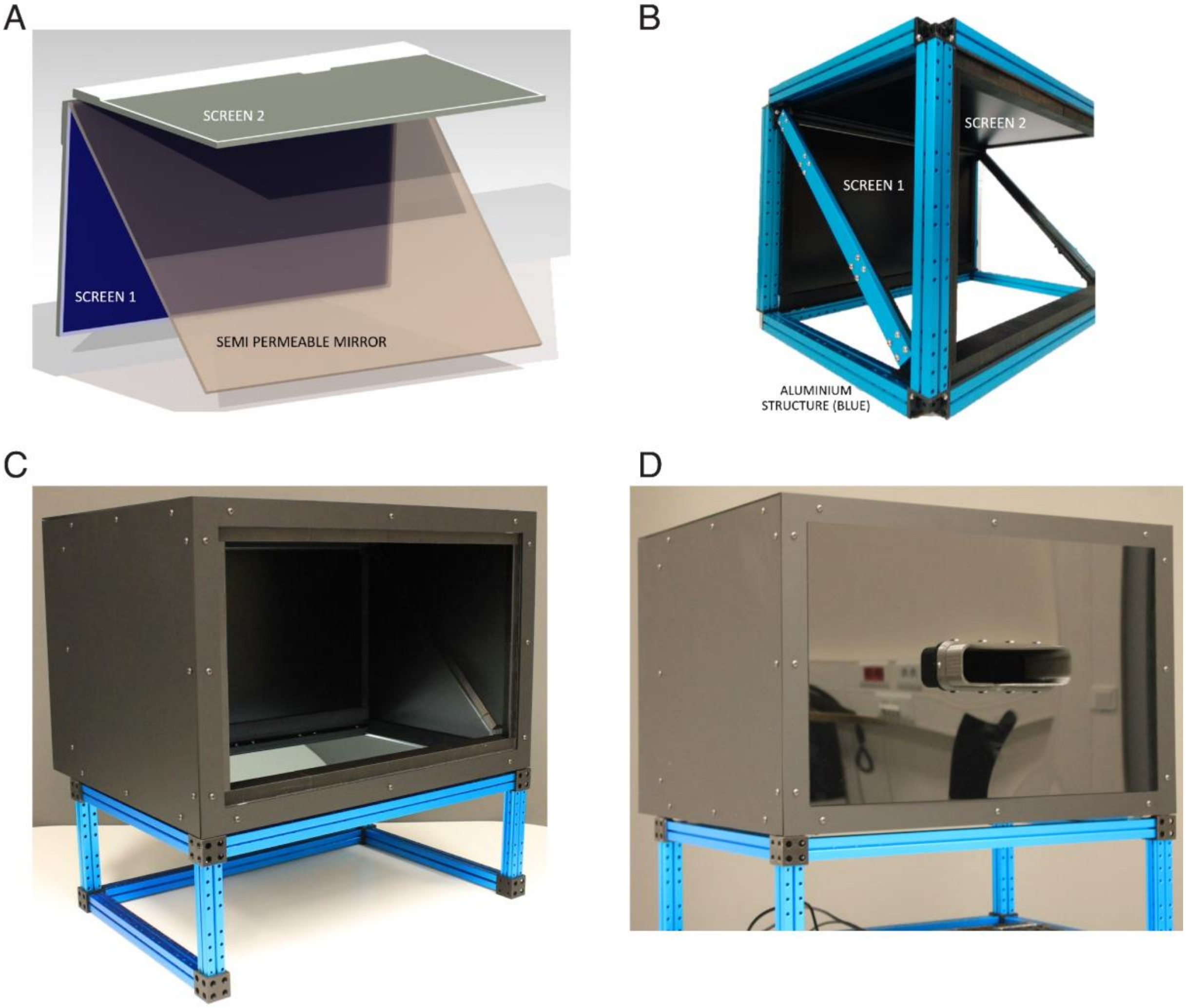


**Supplementary Fig. 1.** LCD Tachistoscope. (A) Vertical screen (screen 1), horizontal screen (screen 2), and semi-permeable mirror. (B) Both screens and the mirror assembled together using an aluminium structure. (C) The tachistoscope assembled without lid and (D) with lid, showing the eyecup through which participants look at the display. Photos taken by Albert De Beir. Reproduced with permission.

**Supplementary Note 2: Face stimuli used in Experiments 1 and 2**

Stimuli were 20 human faces (10 fearful and 10 neutral, the same 10 identities for both, 5 female; **Supplementary Figs. 2-3**) taken from the Radboud Faces Dataset[3] (www.rafd.nl), all seen from a front angle. Images were classified as either fearful or neutral expressions, and were matched between these categories, using ratings from the stimulus dataset, for agreement with the intended emotional expression ($M_{\text{fearful}}$ = 94.9% [$SD_{\text{fearful}}$ = 4.12]; $M_{\text{neutral}}$ = 97.5% [4.03]) while minimising intensity differences (by definition, fearful expressions are higher in intensity than neutral expressions: $M_{\text{fearful}}$ = 4.27 [0.2]; $M_{\text{neutral}}$ = 3.69 [0.27]). Images were cropped to remove hair and create a uniform oval shape, transformed to greyscale, and equated for luminance using the Matlab SHINE toolbox[4]. Scrambled images were created by selecting an oval-shaped area that encompassed all relevant facial features (eyes, nose, and mouth; **Supplementary Fig. 2**). Then, pixels were divided into 40 square patches, which were scrambled by using the Scramble Filter in Photoshop™ CC 2019. The luminance of both intact and scrambled images was equated (mean luminance = 179.8 cd/m$^2$). Background colour was replaced with uniform grey (luminance = 220 cd/m$^2$). Images (2.56° × 3.67° in size) were presented either to the left or to the right of a fixation cross (horizontal centre-to-centre distance 2.82°), between two dots that were placed above and below each stimulus location, serving as placeholders. The vertical distance between placeholder dots was 4.7°.

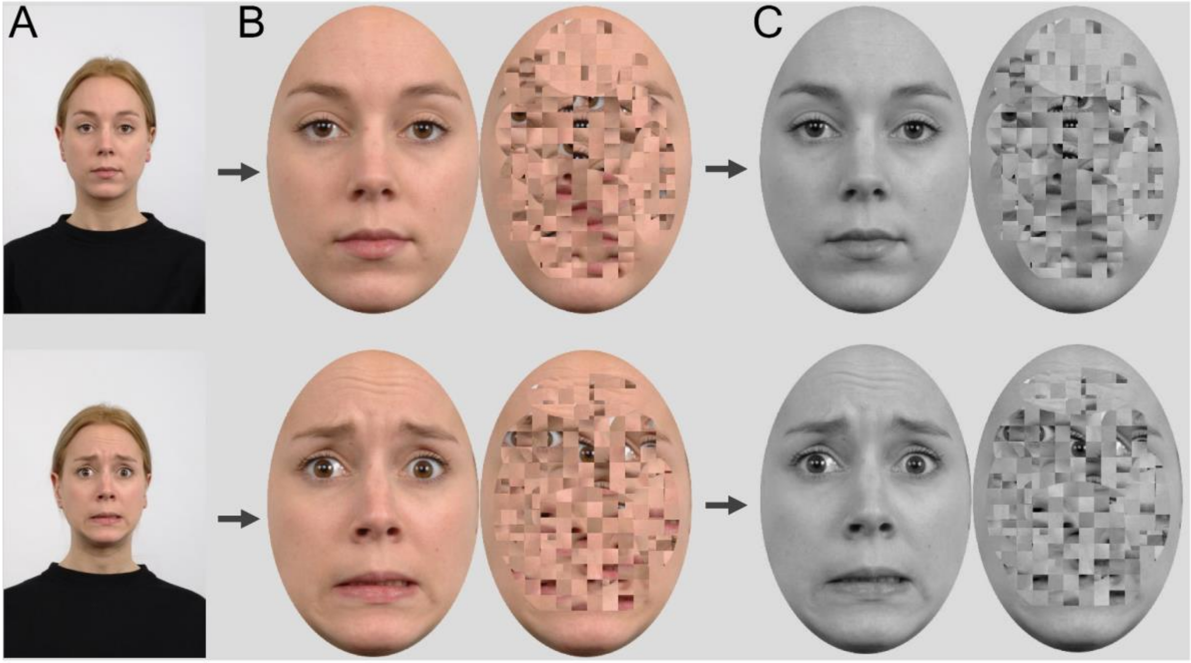


**Supplementary Fig. 2.** Standardisation of face stimuli. (A) Regular stimuli taken from RaFD. (B) Main facial features are preserved; the rest is cropped out. (C) Result of standardisation procedure after equating for luminance and contrast with the Matlab SHINE toolbox. The intact faces shown here belong to the Radboud Face Database (RaFD) and can be presented as stimulus examples (see: https://rafd.socsci.ru.nl/).

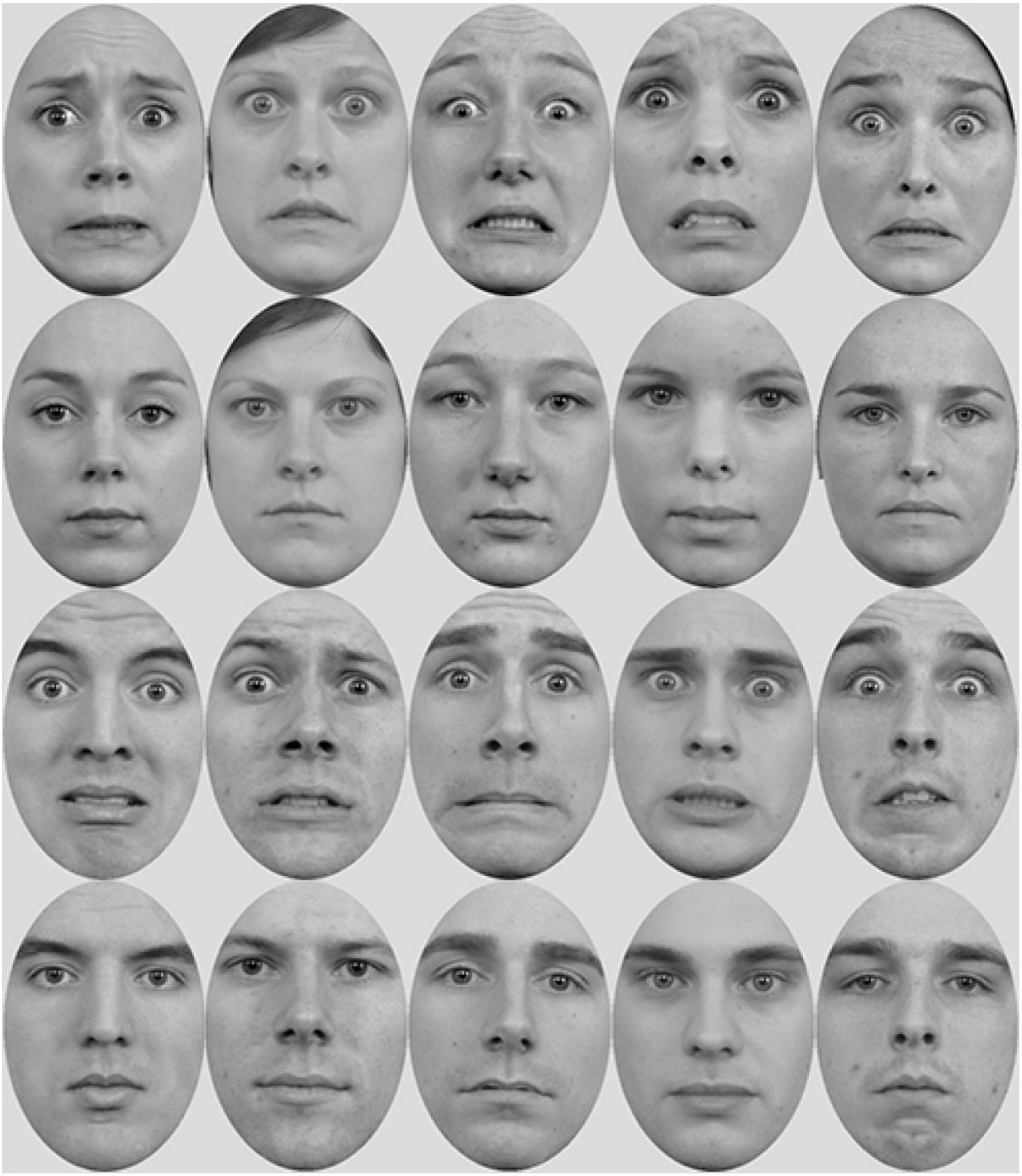

**Supplementary Fig. 3.** RaFD intact faces used in Experiments 1 and 2: fearful and neutral expressions. First row: fearful, female. Second row: neutral, female. Third row: fearful, male. Fourth row: neutral, male. The intact faces shown here belong to the Radboud Face Database (RaFD) and can be presented as stimulus examples (see: https://rafd.socsci.ru.nl/).

All the experiments reported here took place in a dimly lit room. Participants viewed the display through a rectangular eyecup, positioned 55 cm away from the vertical screen. The tachistoscope was connected to a computer running Matlab (version 2018a) and the experiments were written using the Psychophysics Toolbox extensions[5].

### Supplementary Note 3. Full Results of Experiment 1

#### Location sensitivity

To examine how the manipulated factors affected location discrimination, we entered location d′ scores into a 2 (expression: fearful, neutral) × 2 (orientation: upright, inverted) × 7 (exposure durations) repeated-measures ANOVA. We found a main effect of exposure duration, whereby sensitivity increased with increasing exposure duration ($F_{(4.90,\,151.98)} = 215.135, p < .000001$, ηp$^2$ = .874). Crucially, as shown in **Fig. 1B**, location d′ scores went from showing no sensitivity (chance performance) at the shortest exposure duration to showing high sensitivity at the longest exposure durations, indicating that changes in exposure duration in the order of a few milliseconds had a great impact on participants' ability to discriminate an intact face from a scrambled face. Importantly, there was a main effect of face orientation ($F_{(1,\,31)} = 34.918, p = .00000159$, ηp$^2$ = .530), indicating a sensitivity advantage for upright faces ($M = 1.169$ [$SD = 0.905$]) over inverted faces ($M = 0.963$ [0.701]): a face-inversion effect. However, the effect of expression was only marginally significant ($F_{(1,\,31)} = 3.633, p = .066$, ηp$^2$ = .105), with a slight numerical advantage of neutral expressions ($M = 1.088$ [0.977]) over fearful expressions ($M = 1.043$ [0.966]). Importantly, the interaction between face orientation and exposure duration was significant ($F_{(6,\,186)} = 15.331, p < .000001$, ηp$^2$ = .331). To see at what specific exposure duration upright faces enjoyed significantly better sensitivity over inverted faces, we ran post hoc pairwise comparisons, which revealed an advantage of upright faces at 4.4 ms ($t(31) = 5.987, p < .00001, d = 0.671, CI = 0.160 - 0.615$), 5.3 ms ($t(31) = 8.164, p < .000001, d = 0.915, CI = 0.301 - 0.756$), and 6.2 ms of exposure ($t(31) = 5.477, p < .0000128, d = 0.614, CI = 0.127 - 0.582$). Therefore, 4.4 ms of exposure were sufficient to exhibit a face-inversion effect (FIE). We did not find an interaction between expression and face orientation ($F_{(1,\,31)} = 0.001, p = .97$, ηp$^2$ < 0.001), or between expression and exposure duration ($F_{(6,\,186)} = 0.912, p = .487$, ηp$^2$ = .029), or a three-way interaction ($F_{(6,\,186)} = 1.033, p = .405$, ηp$^2$ = .032). Thus, expression had no direct or modulatory effect on location sensitivity.

Although we did not find any effect involving expression, absence of evidence is not necessarily evidence of absence. Therefore, we calculated Bayes factors to test whether the obtained data support the absence of an effect of expression (null hypothesis model). Bayes factors indicated strong evidence in favour of the null hypothesis model ($BF_{01} = 10.571$), suggesting that these data are 10.571 times more likely to be observed under the null hypothesis model of expression. This analysis suggests that fearful expressions are not prioritised for perceptual discrimination when compared to neutral expressions.

Finally, to determine the minimal required exposure for above-chance performance (d′ > 0), we ran a series of uncorrected one-sample t-tests against zero. We found that the earliest exposure duration that elicited above-chance discrimination was 1.7 ms for upright neutral ($M = 0.166$ [0.381]; $t(31) = 2.47$, $p = .019$, $d = 0.436, CI = 0.029 - 0.303$), inverted fearful ($M = 0.228$ [0.396]; $t(31) = 3.25$, $p = .003$, $d = 0.575, CI = 0.085 - 0.370$), and inverted neutral faces ($M = 0.268$ [0.363]; $t(31) = 4.174$, $p = .00022$, $d = 0.738, CI = 0.137 - 0.398$), and 2.6 ms for upright fearful faces ($M = 0.627$ [0.463]; $t(31) = 7.66$, $p < .00001$, $d =$

$1.354, CI = 0.46 - 0.794$). Thus, our results suggest that above-chance discrimination of an intact face stimulus from its scrambled counterpart requires around 2 ms of visual exposure.

### Location response bias

We examined whether participants' response bias for reporting face location varied across conditions by entering the absolute values of $C_{location}$ scores into a 2 (expression: fearful, neutral) × 2 (orientation: upright, inverted) × 7 (exposure durations) repeated-measures ANOVA. Response bias significantly decreased with exposure duration ($F_{(1.73, 53.66)} = 18.692, p < .00001, \eta p^2 = .376$), indicating that as participants' ability to discriminate the face increased (shown by higher location d′ scores) they became less likely to exhibit a systematic bias in their preference to report one side or the other (**Supplementary Fig. 4A**). We did not find a main effect of expression ($F_{(1, 31)} = 0.0004, p = .984, \eta p^2 = .00001$), but we did find an effect of face orientation ($F_{(1, 31)} = 5.59, p = .025, \eta p^2 = .153$), indicating slightly greater bias for inverted faces ($M = 0.454$ [0.177]) over upright faces ($M = 0.388$ [0.210]). To assess whether the obtained data support the absence of an effect of expression, we estimated Bayes factors, which indicated strong evidence for the null hypothesis model ($BF_{01} = 10.380$). No interaction reached significance (all p > .112).

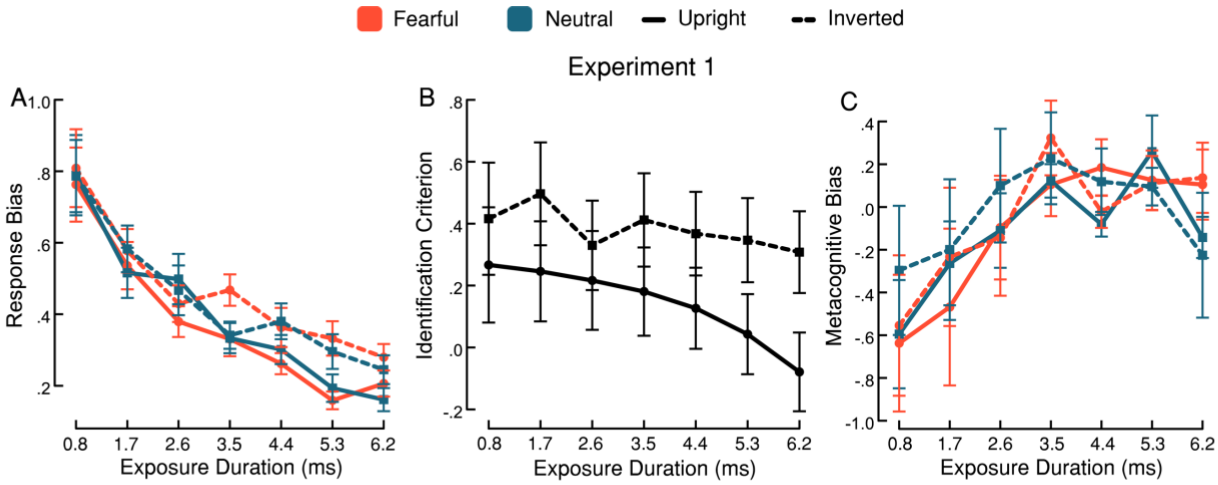


**Supplementary Fig. 4.** Additional results of Experiment 1. (A) Absolute-value response bias scores for reporting location (bias toward either left or right). A repeated-measures ANOVA showed that the amount of response bias decreased as exposure duration increased, with slightly (but significantly) greater bias for inverted faces over upright faces. However, there was no statistically significant difference between fearful and neutral expressions (B) Criterion scores for reporting expression. Although the repeated-measures ANOVA did not find main effects of exposure duration and face orientation, it did find a significant interaction between these two factors. However, Bonferroni-corrected post hoc comparisons did not reveal significant differences. (C) Metacognitive bias scores for reporting subjective awareness. Lower scores denote a more liberal bias to report higher confidence. A repeated-measures ANOVA showed that Meta-bias increased as exposure duration increased, suggesting that participants were more willing to exhibit lower confidence than in shorter exposure durations. Data are presented as mean values with ±1 SEM bars, n = 32 independent participants.

### Expression identification sensitivity

We examined whether participants' sensitivity at identifying emotional expression varied across conditions by entering identification d′ scores – taken over all trials irrespective of location response – into a 2 (orientation: upright, inverted) × 7 (exposure durations) repeated-measures ANOVA (**Fig. 1C**). A main effect of exposure duration indicated that sensitivity to expression increased with increasing exposure duration ($F_{(4.22, 130.93)} = 12.895, p < .00001, \eta p^2 = .294$). We also found a main effect of face orientation ($F_{(1, 31)} = 19.54, p = .0001, \eta p^2 = .387$), such that expression identification d′ was significantly higher for upright faces ($M = 0.243$ [0.235]) than for inverted faces ($M = 0.088$ [0.111]). The interaction between face orientation and exposure duration also reached significance ($F_{(6, 186)} = 3.12, p = .006, \eta p^2 = .091$). To determine the minimal exposure duration that elicited the advantage of upright faces over inverted faces in expression identification, we ran post hoc pairwise comparisons. They revealed a significant advantage of upright faces over inverted faces at 5.3 ms ($t(31) = 3.563,\ p = .041, d = 0.758, CI = 0.004 - 0.54$) and 6.2 ms of exposure ($t(31) = 4.762,\ p = .00033,\ d = 1.013, CI = 0.096 - 0.631$).

To determine the minimal required exposure that exhibited above-chance performance (d′ > 0), we ran a series of uncorrected one-sample t-tests against zero. We found that the earliest exposure duration that elicited above-chance identification was 3.5 ms for upright ($M = 0.220$ [0.349]; $t(31) = 3.57,\ p = .001,\ d = 0.631, CI = 0.094 - 0.346$) and 4.4 ms for inverted faces ($M = 0.137$ [0.291]; $t(31) = 2.67,\ p = .012,\ d = 0.471, CI = 0.032 - 0.242$).

These results indicate that sensitivity to expression identity requires about 5.3 ms of exposure to exhibit a FIE, thereby suggesting that this minimal required exposure was sufficient for the successful integration of facial features in expression identification.

**Expression identification criterion**

We examined whether participants' criteria for reporting fearful expression varied across conditions by entering $C_{identification}$ scores into a 2 (orientation: upright, inverted) × 7 (exposure durations) repeated-measures ANOVA (**Supplementary Fig. 4B**). The lower the value of this measure, the more willing a participant is to report a fearful expression (liberal criterion). We did not find a main effect of exposure duration ($F_{(1.55, 48.11)} = 2.05, p = .150, \eta p^2 = .062$) or of face orientation ($F_{(1, 31)} = 3.11, p = .088, \eta p^2 = .091$). However, we did find a significant interaction between these two factors ($F_{(3.51, 108.73)} = 3.162, p = .021, \eta p^2 = .093$). To determine differences between face orientations per each exposure duration, we used post-hoc pairwise comparisons, but no comparison of relevance reached significance. We calculated Bayes factors to test whether the data support this absence of an orientation effect. Surprisingly, Bayes factors indicated substantial evidence in favour of the alternative hypothesis model ($BF_{01} = 0.0003$), thus suggesting that participants may have exhibited a more liberal criterion when reporting fearful expressions. We also calculated Bayes factors to test whether the data support this absence of an effect of exposure duration. Bayes factors indicated strong evidence in favour of the null hypothesis model ($BF_{01} = 15.959$), confirming that criterion scores did not vary across exposure durations. These results suggest that identification criterion did not become more liberal across increasing exposure duration but might have been more liberal for the identification of fearful faces when the face was presented in an upright orientation than in an inverted orientation.

### Awareness-based metacognitive sensitivity

We examined whether awareness scores were sensitive to participants' location sensitivity scores by estimating meta-d′, a measure of metacognitive sensitivity. To examine whether metacognitive sensitivity varied across conditions, we entered meta-d′ scores into a 2 (expression: fearful, neutral) × 2 (orientation: upright, inverted) × 7 (exposure durations) repeated-measures ANOVA (**Fig. 1D**). A main effect of exposure duration indicated that meta-d′ increased with increasing exposure duration ($F_{(3.69, 114.38)} = 12.922, p < .00001, \eta p^2 = .294$). We also found a main effect of face orientation ($F_{(1, 31)} = 6.475, p = .016, \eta p^2 = .173$), which indicates better metacognitive sensitivity to upright faces ($M = 0.540$ [0.328]) than inverted faces ($M = 0.339$ [0.271]). However, we did not find a main effect of expression ($F_{(1, 31)} = 0.110, p = .742, \eta p^2 = .004$), suggesting that emotional expression did not affect metacognitive sensitivity. We did not find an interaction between face orientation and exposure duration ($F_{(4.11, 127.48)} = 0.512, p = .732, \eta p^2 = .016$), between expression and exposure duration ($F_{(6, 186)} = 0.588, p = .74, \eta p^2 = .019$), between face orientation and expression ($F_{(1, 31)} = 1.048, p = .314, \eta p^2 = .033$), and no three-way interaction either ($F_{(4.48, 138.77)} = 0.321, p = .882, \eta p^2 = .01$). These results suggest upright faces reach conscious access faster than inverted faces.

As described, we did not find a main effect of expression, therefore we calculated Bayes factors to test whether the obtained data support this absence of an effect. Bayes factors indicated strong evidence in favour of the null hypothesis model ($BF_{01} = 13.320$), confirming that neither facial expression is prioritised by metacognitive sensitivity.

Finally, to determine the minimal required exposure that exhibited above-chance metacognitive sensitivity (meta-d′ > 0), we ran a series of uncorrected one-sample t-tests against zero. We found that the earliest exposure duration that elicited above-chance discrimination was 1.7 ms for upright fearful ($M = 0.325$ [0.427]; $t(31) = 4.31, p = .00015, d = 0.762, CI = 0.171 - 0.479$) and upright neutral faces ($M = 0.237$ [0.456]; $t(31) = 2.94, p = .006, d = 0.519, CI = 0.072 - 0.401$), and 2.6 ms for inverted fearful ($M = 0.278$ [0.758]; $t(31) = 2.07$, $p = .047, d = 0.366, CI = 0.005 - 0.551$) and inverted neutral faces ($M = 0.388$ [0.632]; $t(31) = 3.47, p = .002, d = 0.613, CI = 0.16 - 0.616$). Thus, our results suggest that it takes around 2 ms of exposure for a face stimulus to reach above-chance metacognitive sensitivity.

### Metacognitive bias

Metacognitive bias (meta-bias) is the tendency to give high confidence ratings regardless of actual performance. In this experiment, however, we used the PAS, a more exhaustive measure of awareness than confidence ratings[6]. The PAS allows participants to rate their visual experience using four ratings covering from "no experience" to "clear experience" reports. Participants described their visual experience of a face and, therefore, metacognitive bias describes the tendency to describe one's visual experience as clear. To examine whether meta-bias varied across conditions, we entered meta-bias scores into a 2 (expression: fearful, neutral) × 2 (orientation: upright, inverted) × 7 (exposure durations) repeated-measures ANOVA (**Supplementary Fig. 4C**). A main effect of exposure duration indicated that as exposure duration increased, meta-bias became more conservative ($F_{(3.02, 93.65)} = 5.371, p = .002, \eta p^2 = .148$), i.e., participants' tendency to give high confidence ratings decreased with exposure duration. However, we did not find an effect of orientation ($F_{(1, 31)} = 0.749, p = .393, \eta p^2 = .024$) or expression ($F_{(1, 31)} = 0.003, p = .958, \eta p^2 < 0.01$). No interaction reached significance either: between expression and orientation ($F_{(1, 31)} = 0.072, p = .791, \eta p^2 = .002$), expression and exposure duration ($F_{(3.22, 99.88)} = 0.469, p = .718, \eta p^2 = .015$), orientation and exposure duration ($F_{(3.48, 107.84)} = 0.275, p = .870, \eta p^2 = .009$), and the three-way interaction ($F_{(3.41, 105.61)} = 0.299, p = .85, \eta p^2 = .01$).

We calculated Bayes factors to test whether the obtained data support this absence of an effect of orientation and expression. Bayes factors indicated strong evidence in favour of the null hypothesis model of orientation ($BF_{01} = 10.022$) and of the null hypothesis model of expression ($BF_{01} = 13.329$), confirming that meta-bias was unaffected by emotional expression and face orientation.

**Supplementary Note 4: An SDT approach to quantifying perceptual and metacognitive processing**

Our method allows for the quantification of both perceptual sensitivity (the ability to distinguish signal from noise) and metacognitive sensitivity (how participants' subjective experience relates to their perceptual performance). To achieve this, we use response accuracy data from the detection task to calculate d' and response bias (type-1 SDT[7]). Additionally, we combine these data with the Perceptual Awareness Scale (PAS) data to calculate meta-d' and metacognitive bias (type-2 SDT[8]). Importantly, both perceptual and metacognitive sensitivity are bias-independent, meaning they remain unaffected by post-perceptual factors, effectively distinguishing between perceptual and post-perceptual processes. By combining these two SDT approaches, we can quantify the objectively processed sensory information and the portion of that information accessible to awareness (for reviews and discussions, see[9,10]).

In addition to these psychophysical indices, our EEG experiments measured neural indices using univariate and multivariate analyses. This comprehensive approach provides a detailed set of measures for both perceptual processing and awareness of face stimuli and their attributes, enabling direct comparison of the exposure durations required for each measure to arise.

The rationale behind our method for assessing unconscious processing is straightforward: To indicate unconscious processing, the minimal exposure that is required for a marker of perceptual processing to arise should be shorter than the minimal required exposure for markers of awareness.

The rationale for our method and conclusions aligns with the logic of previous investigations that interpret processing in the absence of awareness as evidence for unconscious processing[6,11–13]. Most of the studies providing evidence for unconscious holistic processing[14–23] and unconscious emotion processing[24–32] of faces assessed awareness using dichotomous ratings (but see[33]). In behavioural masking studies, the masks ensure the invisibility of the stimuli of interest and processing is typically measured with a spatial detection task; the logic suggesting that higher-than-chance performance in this task (with successful suppression from awareness) indicates unconscious processing is similar to our own analyses' rationale, but the procedure lacks the fine control of exposure that our method affords. In interocular suppression studies, such as those using the breaking continuous flash suppression method, differences in the time it takes for suppressed stimuli to break through interocular suppression are interpreted as evidence for unconscious processing. This method makes several assumptions about the relationship between reaction times and awareness; those assumptions have been criticised, calling into question the validity of unconscious processing claims based on this method[20,21,34–39] and raising the need to use more rigorous methods to make such inferences.

The extreme temporal precision of our visual displays, along with the multitude of behavioural and neural markers, enables extreme sensitivity to the possible presence of unconscious processing. Moreover, the richness of our data sets a high standard for drawing conclusions about the relationship between perception and awareness, as a range of markers of perceptual processing and awareness must converge to establish convincing claims. For example, if markers of holistic face processing (e.g., FIE, N170/VPP, MVPA) were to arise with similarly shorter exposure durations as those required for awareness, this would strongly support the existence of unconscious face processing.

**Supplementary Note 5. Full Results of Experiment 2**

**Order sensitivity**

To examine how the manipulated factors affected face discrimination, we entered order d′ scores into a 2 (expression: fearful, neutral) × 2 (orientation: upright, inverted) × 7 (exposure durations) repeated-measures ANOVA. We found a main effect of exposure duration, whereby sensitivity increased with increasing exposure duration ($F_{(2.23,\,69.02)} = 180.786, p < .00001, \eta p^2 = .854$). As in Experiment 1 with location d′ scores, here order d′ scores went from showing no sensitivity (chance performance) at the shortest exposure duration to showing high sensitivity at the longest exposure durations (**Fig. 1E**). We also found a main effect of face orientation ($F_{(1,\,31)} = 49.058, p < .00001, \eta p^2 = .613$), indicating a sensitivity advantage for upright faces ($M = 0.970$ [0.785]) over inverted faces ($M = 0.709$ [0.616]), but did not find a main effect of expression ($F_{(1,\,31)} = 0.761, p = .39, \eta p^2 = .024$), indicating no advantage of fearful expressions over neutral expressions. Importantly, the interaction between face orientation and exposure duration was significant ($F_{(6,\,186)} = 10.713, p < .00001, \eta p^2 = .257$). To see at what specific exposure durations upright faces enjoyed significantly better sensitivity over inverted faces, and thus answer our main question, we ran post hoc Bonferroni-corrected pairwise comparisons, which revealed said advantage at 3.3 ms ($t(31) = 4.737$, $p = .00039$, $d = 0.584, \mathrm{CI} = 0.086 - 0.578$), 4.2 ms ($\mathrm{t}(31) = 7.515$, $\mathrm{p} < .00001$, $\mathrm{d} = 0.927, \mathrm{CI} = 0.28 - 0.773$), 5.1 ms ($\mathrm{t}(31) = 6.694$, $\mathrm{p} < .00001$, $\mathrm{d} = 0.825, \mathrm{CI} = 0.223 - 0.715$), and 6 ms of exposure ($\mathrm{t}(31) = 4.837$, $\mathrm{p} = .00025$, $\mathrm{d} = 0.596, \mathrm{CI} = 0.093 - 0.585$). Crucially, 3.3 ms of exposure were sufficient to elicit a face-inversion effect when perceptual discrimination relied on foveal vision. As in Experiment 1, we did not find an interaction between expression and face orientation ($\mathrm{F}_{(1,\,31)} = 0.504, \mathrm{p} = .483, \eta p^2 = .016$), or between expression and exposure duration ($\mathrm{F}_{(6,\,186)} = 0.328, \mathrm{p} = .899, \eta p^2 = .01$), or a three-way interaction ($\mathrm{F}_{(6,\,186)} = 1.313, \mathrm{p} = .253, \eta p^2 = .041$).

To determine the minimal exposure that exhibited above-chance performance (d′ > 0), we ran a series of uncorrected one-sample t-tests against zero. We found that the earliest exposure duration that elicited above-chance discrimination was 1.5 ms for upright fearful ($M = 0.197$ [0.299]; $t(31) = 3.72, p < .00001, d = 0.658, \mathrm{CI} = 0.089 - 0.305$), and 2.4 ms for upright neutral faces ($M = 0.450$ [0.534]; $t(31) = 4.77, p < .00001, d = 0.842, \mathrm{CI} = 0.257 - 0.642$), inverted fearful faces ($M = 0.358$ [0.364]; $t(31) = 5.57, p < .00001, d = 0.984, \mathrm{CI} = 0.227 - 0.490$), and inverted fearful faces ($M = 0.367$ [0.534]; $t(31) = 3.90, p = .00048, d = 0.689, \mathrm{CI} = 0.175 - 0.560$). Our results suggest that it takes around 2 ms of exposure for a face stimulus to reach above-chance discrimination from its scrambled counterpart when relying on foveal vision.

Like in Experiment 1, we did not find a main effect of expression. Therefore, we calculated Bayes factors to test whether the obtained data support the absence of this effect (null hypothesis model). Bayes factors indicated strong evidence in favour of the null hypothesis model ($BF_{01}$ = 11.891). This analysis suggests that fearful expressions are not prioritised by perceptual sensitivity over neutral expressions.

### Order response bias

We examined whether participants' response bias for reporting the intact face presentation order varied across conditions by entering the absolute values of $C_{order}$ scores into a 2 (expression: fearful, neutral) × 2 (orientation: upright, inverted) × 7 (exposure durations) repeated-measures ANOVA. As in Experiment 1, response bias significantly decreased with increasing exposure duration ($F_{(1.98, 61.31)} = 12.581, p < .00001, \eta p^2 = .289$), indicating that as participants' ability to discriminate the face increased, they became less likely to exhibit a systematic bias in their preference to report one presentation order or the other (**Supplementary Fig. 5A**). We did not find a main effect of expression ($F_{(1, 31)} = 0.014, p = .907, \eta p^2 < 0.01$), or of face orientation ($F_{(1, 31)} = 1.099, p = .303, \eta p^2 = .034$). No interaction reached significance (all p > .261).

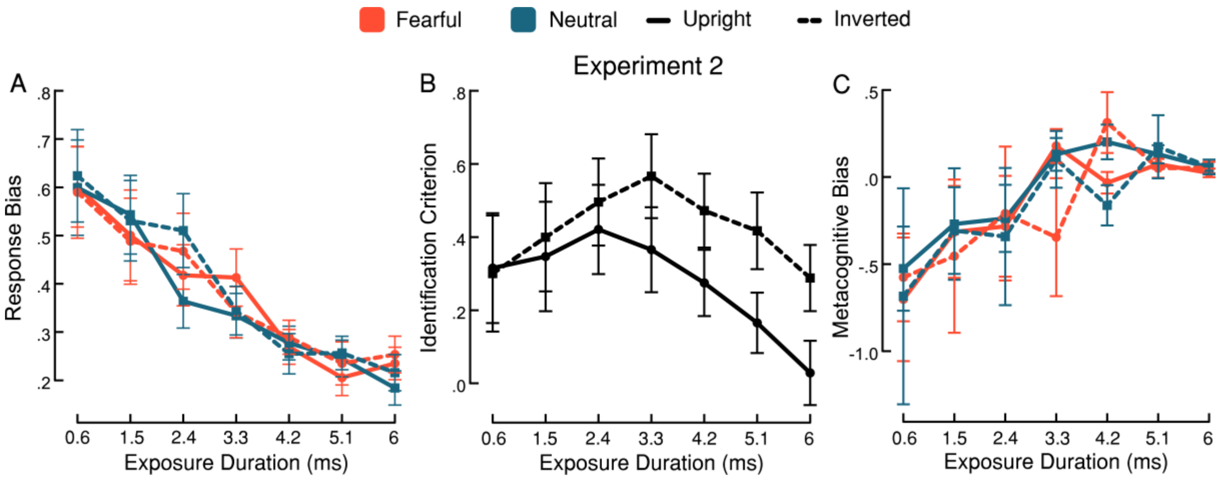


**Supplementary Fig. 5.** (A-C) Additional results of Experiment 2. (A) Absolute-value response bias scores for reporting presentation order of the intact face (bias toward reporting either first or second). A repeated-measures ANOVA showed that the amount of bias decreased as exposure duration increased, but there was no difference in amount of response bias between expressions and orientations. (B) Criterion scores for reporting expression. A repeated-measures ANOVA showed that upright faces exhibit a significantly more liberal criterion (lower values) than inverted faces during expression identification. (C) Metacognitive bias scores for reporting subjective awareness. Lower scores denote a more liberal bias to report higher confidence. A repeated-measures ANOVA showed that meta-bias increased as exposure duration increased, suggesting that participants were more willing to exhibit lower confidence than in shorter exposure durations. Data are presented as mean values with ±1 SEM bars, n = 32 independent participants.

To assess whether the obtained data support the absence of an effect of expression and of orientation, we estimated Bayes factors, which indicated strong evidence for the null hypothesis model of expression ($BF_{01}$ = 13.746) and of orientation ($BF_{01}$ = 10.287).

### Expression identification sensitivity

We examined whether participants' sensitivity at identifying expression varied across conditions by entering identification d′ scores into a 2 (orientation: upright, inverted) × 7 (exposure durations) repeated-measures ANOVA (**Fig. 1F**). We found a main effect of exposure duration, indicating that sensitivity to expression increased with increasing exposure duration ($F_{(3.36,\,104.12)} = 36.20, p < .00001, \eta p^2 = .539$). We also found a main effect of face orientation ($F_{(1,\,31)} = 27.867, p = .0000097, \eta p^2 = .473$), such that expression identification sensitivity was significantly higher for upright faces ($M$ = 0.389 [0.418]) than for inverted faces ($M$ = 0.155 [0.227]). The interaction between orientation and exposure duration also reached significance ($F_{(6,\,186)} = 6.077, p < .00001, \eta p^2 = .164$). To determine the minimal exposure duration that elicited this face-inversion effect in expression identification sensitivity, we ran post hoc Bonferroni-corrected pairwise comparisons. They revealed significant advantages of upright faces over inverted faces at 4.2 ms ($t(31) = 3.967, p = .009, d = 0.843, \mathrm{CI} = 0.04 - 0.657$), 5.1 ms ($t(31) = 4.632$, $p = .00059$, $d = 0.984, \mathrm{CI} = 0.098 - 0.716$), and 6 ms of exposure ($t(31) = 6.279, p < .00001, d = 1.334, \mathrm{CI} = 0.243 - 0.861$).

To determine the minimal exposure that exhibited above-chance performance (d′ > 0) in expression identification, we ran a series of uncorrected one-sample t-tests against zero. We found that the earliest exposure duration that elicited above-chance discrimination was 1.5 ms for upright faces ($M$ = 0.112 [0.262]; $t(31) = 2.41$, $p = .022$, $d = 0.427, \mathrm{CI} = 0.017 - 0.207$), and 3.3 ms for inverted faces ($M$ = 0.169 [0.322]; $t(31) = 2.96$, $p = .006$, $d = 0.524, \mathrm{CI} = 0.053 - 0.285$).

#### Expression identification criterion

We examined whether participants' criterion for reporting fearful expression varied across conditions by entering $C_{identification}$ scores into a 2 (orientation: upright, inverted) × 7 (exposure durations) repeated-measures ANOVA. We did not find a significant main effect of exposure duration ($F_{(1.46,\,45.36)} = 2.66, p = .096, \eta p^2 = .079$), but we did find a main effect of face orientation ($F_{(1,\,31)} = 8.91, p = .005, \eta p^2 = .223$), indicating a more liberal criterion for upright faces ($M$ = 0.274 [0.135]) than inverted faces ($M$ = 0.420 [0.102]), i.e., participants were more willing to report a fearful expression for an upright face than for an inverted face (**Supplementary Fig. 5B**). The interaction between face orientation and exposure duration reached significance ($F_{(3.79,\,117.43)} = 3.26, p = .016, \eta p^2 = .095$). To determine differences between face orientations per each exposure duration, we ran post hoc Bonferroni-corrected pairwise comparisons. However, no comparison of interest reached significance. To assess whether the obtained data support the absence of an effect of exposure duration, we estimated Bayes factors, which unexpectedly indicated anecdotal evidence for the alternative hypothesis model ($BF_{01}$ = 0.046), i.e., in favour of an effect of exposure duration. These results suggest that identification criterion was only affected by face orientation. However, the data are inconclusive regarding whether exposure duration contributed to identification criterion.

#### Awareness-based metacognitive sensitivity

We examined whether awareness scores were sensitive to participants' order sensitivity scores by estimating meta-d′. To examine whether metacognitive sensitivity varied across conditions, we entered meta-d′ scores into a 2 (expression: fearful, neutral) × 2 (orientation: upright, inverted) × 7 (exposure durations) repeated-measures ANOVA (**Fig. 1G**). A main effect of exposure duration indicated that meta-d′ increased with increasing exposure duration ($F_{(3.41,\,105.73)} = 29.216, p < .00001, \eta p^2 = .485$). We also found a main effect of face orientation ($F_{(1,\,31)} = 6.176, p = .019, \eta p^2 = .166$), which indicates better metacognitive sensitivity to upright faces ($M = 0.573$ [0.461]) than inverted faces ($M = 0.440$ [0.370]). However, we did not find a main effect of expression ($F_{(1,\,31)} < 0.01, p = .977, \eta p^2 < 0.01$), thus suggesting emotional expression did not affect metacognitive sensitivity. No other interaction reached significance (all $p > .298$). These results suggest that upright faces reach awareness faster than inverted faces.

As described, we did not find a main effect of expression, therefore we calculated Bayes factors to test whether the obtained data support this absence of an effect. Bayes factors indicated strong evidence in favour of the null hypothesis model ($BF_{01} = 13.343$), thus suggesting that fearful expressions are not prioritised by metacognitive sensitivity when compared to neutral expressions.

Finally, to determine the minimal exposure that exhibited above-chance metacognitive sensitivity (meta-d′ > 0), we ran a series of uncorrected one-sample t-tests against zero. We found that the earliest exposure duration that elicited above-chance discrimination was 2.4 ms for upright fearful faces ($M = 0.374$ [0.587]; $t(31) = 3.60, p = .001, d = 0.637, \mathrm{CI} = 0.162 - 0.586$), upright neutral faces ($M = 0.307$ [0.597]; $t(31) = 2.91, p = .007, d = 0.514, \mathrm{CI} = 0.092 - 0.522$), inverted fearful faces ($M = 0.273$ [0.706]; $t(31) = 2.18, p = .037, d = 0.386, \mathrm{CI} = 0.018 - 0.527$), and 3.3 ms for inverted neutral faces ($M = 0.507$ [0.535]; $t(31) = 5.36, p = .000076, d = 0.948, \mathrm{CI} = 0.314 - 0.7$). Thus, our results suggest that it takes ~3 ms of exposure for a face stimulus to reach above-chance metacognitive sensitivity.

**Metacognitive bias**

We estimated meta-bias as in Experiment 1. To examine whether meta-bias varied across conditions, we entered meta-bias scores into a 2 (expression: fearful, neutral) × 2 (orientation: upright, inverted) × 7 (exposure durations) repeated-measures ANOVA (**Supplementary Fig. 5C**). A main effect of exposure duration indicated that meta-bias became less liberal with increasing exposure duration ($F_{(3.21,\,99.55)} = 4.127, p = .007, \eta p^2 = .117$), thus indicating that as participants' metacognitive sensitivity increased, they became less likely to systematically report having no experience (or clear experience) of the intact stimulus shown. However, we did not find an effect of face orientation ($F_{(1,\,31)} = 0.420, p = .522, \eta p^2 = .013$) or of expression ($F_{(1,\,31)} = 0.139, p = .712, \eta p^2 = .004$). No interaction reached significance either (all $p > .550$).

We calculated Bayes factors to test whether the obtained data support this absence of an effect of orientation and expression. Bayes factors indicated strong evidence in favour of the null hypothesis model of orientation ($BF_{01} = 11.326$) and of expression ($BF_{01} = 12.287$), thus suggesting that neither fearful (compared to neutral) expressions nor upright (compared to inverted) faces are prioritised by metacognitive sensitivity.

**Supplementary Note 6: Control experiments**

**Control Experiment 1**

To avoid creating a positive afterimage (i.e., an afterimage of the afterimage-like stimuli), we used backward masking. In addition, we only used one exposure duration of 10 ms, an exposure duration that should be sufficiently long to elicit the FIE both with location sensitivity scores and with expression identification sensitivity scores, as shown in Experiment 1. If afterimage processing made a substantial contribution to the findings reported in Experiment 1, we should find these two FIEs with these new afterimage-like stimuli. If we do not, these findings in Experiment 1 cannot be attributed to afterimage processing.

**Participants**

All the participants of Experiment 1 did this control experiment 10 minutes after having finished Experiment 1.

**Stimuli**

Stimuli were the same as in Experiment 1, but with inverted colours to emulate negative afterimages, including both intact and scrambled face images (**Supplementary Fig. 6**); see **Supplementary Fig. 7** for a full presentation of the afterimage-like stimuli used in the two control experiments.

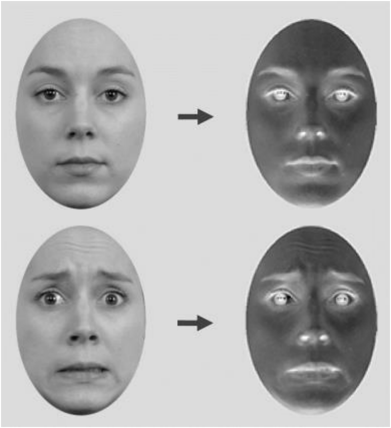

**Supplementary Fig. 6.** Examples of afterimage-like stimuli. (Left) Intact face with neutral (top) and fearful (bottom) expression. (Right) Their colour-inverted afterimage-like counterparts. The intact faces shown here belong to the Radboud Face Database (RaFD) and can be presented as stimulus examples (see: https://rafd.socsci.ru.nl/).

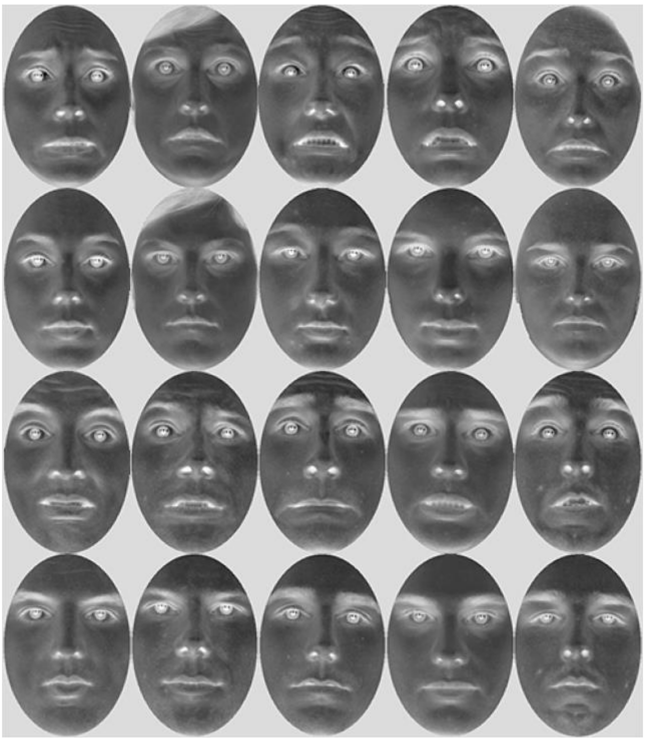

**Supplementary Fig. 7.** Stimuli used in Control Experiments 1 and 2. First row: fearful expressions, female. Second row: neutral expressions, female. Third row: fearful expressions, male. Fourth row: neutral expressions, male. The original faces shown here belong to the Radboud Face Database (RaFD) and can be presented as stimulus examples (see: https://rafd.socsci.ru.nl/).

### Procedure

The procedure was similar to that of Experiment 1, but with only one exposure duration (10 ms); hence this experiment had fewer trials in total (160). In addition, backward masking was used to prevent the new stimuli from creating afterimages. To achieve this, on each trial we presented the non-colour-inverted scrambled version of the face used in that trial. The mask was displayed for 50 ms immediately after stimulus offset in both intact and scrambled stimulus locations. Then, as in Experiment 1, participants were asked to first judge the location and expression of the intact face stimulus, followed by rating their visual experience (PAS).

### Results

#### Location sensitivity

To examine whether face orientation and expression modulate participants' ability to discriminate the location of afterimage-like face stimuli, we entered location d′ scores into a 2 (expression: fearful, neutral) × 2 (orientation: upright, inverted) repeated-measures ANOVA (**Supplementary Fig. 8A**). We did not find an effect of expression ($F_{(1, 31)} = 0.138, p = .713, \eta p^2 = .004$) or of orientation ($F_{(1, 31)} = 0.393, p = .535, \eta p^2 = .013$), indicating that neither factor modulated location d′. The interaction between these two factors did not reach significance either ($F_{(1, 31)} = 3.267, p = .08, \eta p^2 = .095$), even though stimuli were presented for substantially longer than the duration that enabled such discriminations for intact faces. To assess whether the obtained data support the absence of an effect of expression and of orientation, we estimated Bayes factors, which indicated substantial evidence for the null hypothesis model in the former ($BF_{01} = 5.153$) and in the latter ($BF_{01} = 4.974$).

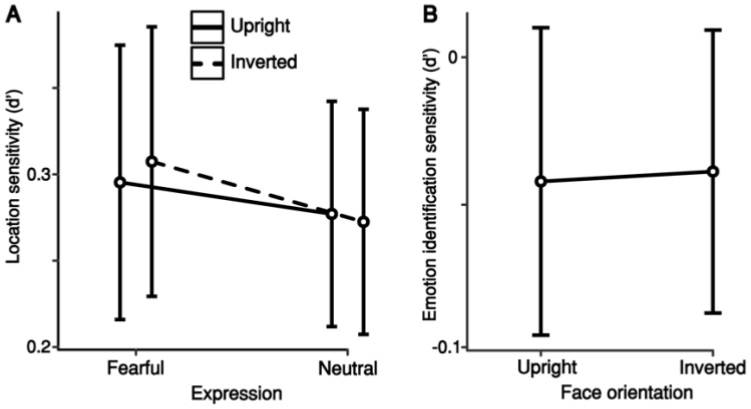


**Supplementary Fig. 8.** Results of Control Experiment 1. (A) Location sensitivity. A repeated-measures ANOVA showed that location d′ scores were very low and unaffected by expression or orientation. (B) Identification sensitivity for expression. A repeated-measures ANOVA showed that emotion identification d′ was at chance level both for upright and inverted faces. Data are presented as mean values with ±1 SEM bars, n = 32 independent participants.

To examine whether participants could tell, above chance, on what side the intact afterimage-like faces were, we ran a series of uncorrected one-sample t-tests against zero. We found that upright fearful ($M = 0.264$ [0.374]; $t(31) = 4.66,\ p < .00001, d = 0.824, CI = 0.167 - 0.426$), upright neutral ($M = 0.280$ [0.306]; $t(31) = 5.4,\ p < .00001, d = 0.954, CI = 0.175 - 0.388$), inverted fearful ($M = 0.253$ [0.382]; $t(31) = 4.9,\ p < .00001, d = 0.866, CI = 0.179 - 0.433$), and inverted neutral faces ($M = 0.263$ [0.305]; $t(31) = 5.33,\ p < .00001, d = 0.943, CI = 0.172 - 0.384$) were discriminated above chance, thus indicating that participants could discriminate between intact and scrambled afterimage-like faces, though with low sensitivity.

#### Expression identification sensitivity

To examine whether face orientation modulates participants′ ability to identify the expression of afterimage-like face stimuli, we compared identification d′ scores for upright and inverted faces by using a paired-sample t-test. We did not find a significant difference between face orientations in identification sensitivity ($t(31) = 0.808, p = .425, d = 0.143, \mathrm{CI} = -0.01 - 0.004$). To assess whether the obtained data support the absence of an effect of face orientation, we estimated Bayes factors, which indicated substantial evidence for the null hypothesis model ($BF_{01}$ = 3.920). These results suggest that participants are not able to identify fearful expressions and distinguish them from neutral expressions by using afterimage processing (**Supplementary Fig. 8B**).

#### Control Experiment 2

In Control Experiment 1, we tested whether findings in Experiment 1 could be affected by afterimage processing. Our findings suggested that neither face orientation nor expression could be discriminated with afterimage-like stimuli presented for 10 ms (over 3 ms longer than the longest-duration stimuli in Experiment 1). However, that approach presents two limitations. First, we used backward masking to prevent those afterimage-like stimuli from creating additional afterimages. Yet backward masking could have also disrupted perceptual sensitivity, thus making the procedure less reliable. Second, we only used one exposure duration, making it impossible to determine whether exposure duration modulates the discrimination of intact afterimage-like stimuli from their scrambled afterimage-like counterparts. In this second control experiment, we addressed these issues with the 2IFC task used in Experiment 2.

We employed exposure durations that covered the range of values used in Experiment 2: from when order sensitivity was at chance (0.6 ms*)*, when the FIE arose (3.3 ms), and when location sensitivity was very high (6 ms). We used the same afterimage-like stimuli of Control Experiment 1, but this time we did not use any masking. If these stimuli exhibit a modulatory effect of face orientation on order sensitivity (a FIE), it is likely that afterimage processing may have contributed to these same effects in Experiment 2. Conversely, if these stimuli do not exhibit said effect, it would suggest that the FIE found in Experiment 2 cannot be explained by afterimage processing.

#### Stimuli, Procedure, and Analyses

Stimuli were the same as in Control Experiment 1, and procedures were the same as in Experiment 2, but with only three exposure durations, hence fewer trials in total (480). Analyses were the same as in Experiment 2 for sensitivity and criterion.

#### Results

#### Order sensitivity

To examine how the manipulated factors affected order presentation discrimination of afterimage-like faces, we entered order d′ scores into a 2 (expression: fearful, neutral) × 2 (orientation: upright, inverted) × 3 (exposure durations) repeated-measures ANOVA. Importantly, a main effect of exposure duration was found ($F_{(2, 62)} = 114.565, p < .00001, \eta p^2 = .787$) indicating that order sensitivity increased with increasing exposure duration when using afterimage-like faces (**Supplementary Fig. 9A**). However, we did not find an effect of expression ($F_{(1, 31)} = 0.011, p = .917, \eta p^2, 0.01$) nor of orientation ($F_{(1, 31)} = 2.698, p = .111, \eta p^2 = .080$), thus suggesting neither expression nor face orientation affected sensitivity to afterimage-like faces. Importantly, this finding of a null effect of orientation suggests that the orientation effect found in Experiment 2 cannot be attributed to afterimage processing. The interaction between face orientation and exposure duration reached significance ($F_{(2, 62)} = 4.228, p = .019, \eta p^2 = .12$). However, post hoc Bonferroni-corrected pairwise comparisons did not reveal significant differences between upright and inverted faces at any exposure duration. No other interaction reached significance (all p > .458). To assess whether the obtained data support the absence of an effect of expression and of face orientation, we estimated Bayes factors, which indicated substantial evidence for the null hypothesis model in the former ($BF_{01} = 8.394$) and in the latter ($BF_{01} = 5.849$). These results suggest that discrimination of afterimage-like faces increases with exposure duration similarly as it did with regular images in Experiment 2. However, face orientation did not modulate discrimination this time, which may suggest that either afterimage processing does not contribute to holistic face processing or that it does but requires longer exposure durations to reveal a FIE.

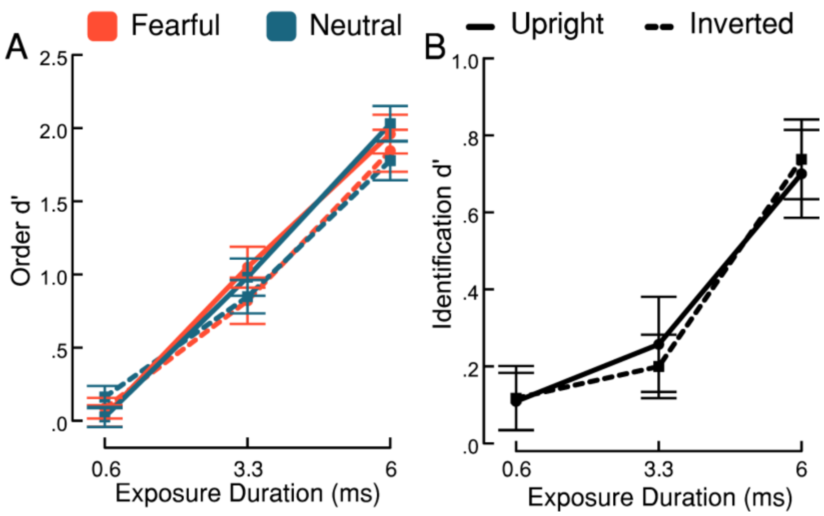


**Supplementary Fig. 9**. Results of Control Experiment 2. (A) Order sensitivity. A repeated-measures ANOVA showed that order d′ increased with increasing exposure duration from chance discrimination to high discrimination but was unaffected by expression or orientation. (B) Identification sensitivity for expression. A repeated-measures ANOVA showed that identification d′ increased with increasing exposure duration but was unaffected by orientation. Data are presented as mean values with ±1 SEM bars, n = 32 independent participants.

### Expression identification sensitivity

To examine how the manipulated factors affected expression identification of afterimage-like faces, we compared identification d′ scores for upright and inverted faces by entering the data into a 2 (orientation: upright, inverted) × 3 (exposure durations) repeated-measures ANOVA. A main effect of exposure duration was found ($F_{(2, 62)} = 16.521, p = .0000017, \eta p^2 = .348$), suggesting that expression identification sensitivity increased with increasing exposure duration (**Supplementary Fig. 9B**). However, we did not find an effect of face orientation ($F_{(1, 31)} = 0.004, p = .952, \eta p^2 < 0.01$), indicating that face orientation did not affect expression identification. Likewise, the interaction between these factors did not reach significance ($F_{(2, 62)} = 0.195, p = .824, \eta p^2 = .006$). To assess whether the obtained data support the absence of an effect of face orientation, we estimated Bayes factors, which indicated substantial evidence for the null hypothesis model ($BF_{01} = 6.648$). These results suggest that expression identification increases with increasing exposure duration regardless of face orientation. Because turning faces upside down disrupts holistic face processing, these results might rely on discrimination of local face features preserved in afterimage-like faces rather than on holistic features.

**Discussion of control experiments**

When a visual image is presented and then removed, retinal impressions (afterimages) can be formed. Because we could not measure, in Experiments 1 and 2, whether such afterimages were formed, to what extent they were processed, and whether they contributed to participants' sensitivity, we ran two control experiments to assess participants' sensitivity to afterimage-like images of faces and assess whether the results of Experiments 1 and 2 could be attributed – in whole or in part – to afterimage processing. In Control Experiments 1 and 2, we used afterimage-like faces to emulate afterimage processing by measuring participants' sensitivity to relatively long presentations of those stimuli (compared to the durations used in Experiment 1), and backward masking to prevent said afterimage-like stimuli from creating additional afterimages. We found above-chance (though weak) location sensitivity; participants were unable to identify expressions, though, and the inversion effect seen in Experiment 1 was absent. However, our procedure in Control Experiment 1 was different from that of Experiment 1 – our use of backward masking may have interacted with stimulus processing, and the use of a single presentation duration did not allow us to measure whether exposure duration modulated perceptual sensitivity.

In Control Experiment 2, we presented participants with unmasked stimuli, using three of the exposure durations used in Experiment 2. We found that participants could discriminate intact afterimage-like faces from their scrambled counterparts as exposure duration increased. Crucially, they did not exhibit a FIE either in order discrimination or expression identification. These results suggest that in both Experiments 1 and 2 participants' sensitivity to faces might have partially relied on information coming from afterimage processing, but it is very unlikely that such information contributed to holistic face processing as indexed by the FIE or to expression identification.

**Supplementary Note 7. Additional details of Experiment 3**

Detecting a single stimulus can be performed using low-level stimulus attributes (without extracting meaning), reducing the exposure required for localisation. Therefore, we took advantage of our LCD tachistoscope's ability to display stimuli for sub-millisecond durations and selected seven new equally-spaced durations (range 0.25 – 1.25 ms) covering floor to ceiling localisation performance. To choose these durations, we ran a pilot study where six participants detected the location of neutral upright faces presented for a range of very short durations (0.1–1 ms in 0.05 ms increments, and 1–1.6 ms in 0.1 ms increments; 20 trials/duration). We used the results shown in **Supplementary Fig. 10** to choose seven new equally-spaced presentation durations for Experiment 3, capturing the range of performance from floor to ceiling (0.250, 0.417, 0.583, 0.750, 0.917, 1.083, and 1.250 ms of exposure).

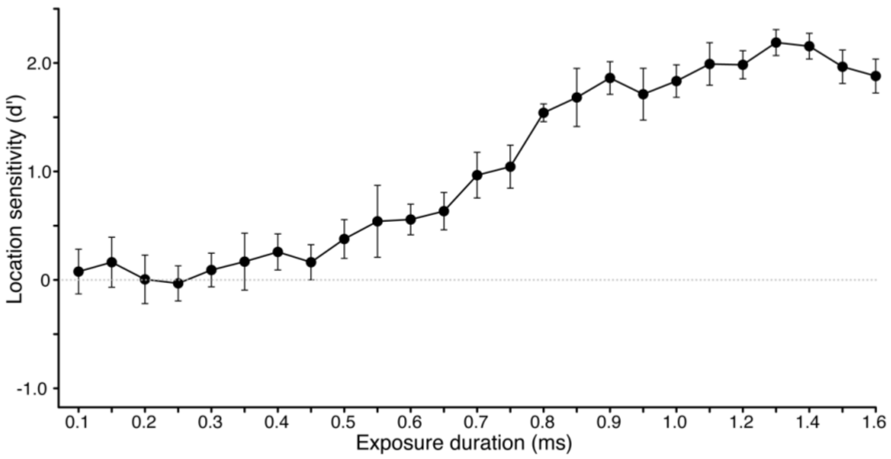


**Supplementary Fig. 10**. Pilot results for Experiment 3. Location sensitivity increased from chance level to high sensitivity across exposure durations. Data are presented as mean values with ±1 SEM bars, n = 6 independent participants.

### Location sensitivity

To examine how the manipulated factors affected stimulation location detection, we entered location d′ scores into a 2 (expression: fearful, neutral) × 2 (orientation: upright, inverted) × 7 (exposure durations) repeated-measures ANOVA. We found a main effect of exposure duration ($F_{(2.68,\,83.19)} = 300.801, p < .00001, \eta p^2 = .907$). As shown in **Fig. 2A**, location d′ scores increased from no sensitivity at the shortest exposure duration to high sensitivity at the longest exposure durations, indicating that changes in exposure duration between 0.25 and 1.25 ms had great impact on participants' ability to detect stimulation. We also found a main effect of face orientation ($F_{(1, 31)} = 4.69, p = .038, \eta p^2 = .131$). We did not find a main effect of expression ($F_{(1, 31)} = 2.031, p = .164, \eta p^2 = .061$). No interaction reached significance: While the interaction between exposure duration and face orientation was marginally significant ($F_{(6, 186)} = 2.009, p = .067, \eta p^2 = .061$), the other interactions were not (all $p > .303$).

#### Location response bias

We examined whether participants' response bias for reporting the face presentation location varied across conditions by entering the absolute values of $C_{order}$ scores into a 2 (expression: fearful, neutral) × 2 (orientation: upright, inverted) × 7 (exposure durations) repeated-measures ANOVA. Response bias significantly decreased with increasing exposure duration ($F_{(1.7, 52.55)} = 16.631, p = .0000079, \eta p^2 = .349$), indicating that as participants' ability to detect stimulation increased, they became less likely to exhibit a systematic bias in their preference to report one presentation location or the other (**Supplementary Fig. 11A**). We did not find a main effect of expression ($F_{(1, 31)} = 0.64, p = .43, \eta p^2 = 0.02$), or of face orientation ($F_{(1, 31)} = 0.562, p = .459, \eta p^2 = .018$). The interaction between exposure duration and expression reached significance ($F_{(6, 186)} = 2.59, p = .02, \eta p^2 = 0.08$), but no post-hoc pairwise comparison reached significance (all $p > .095$). No other interaction reached significance (all $p > .293$).

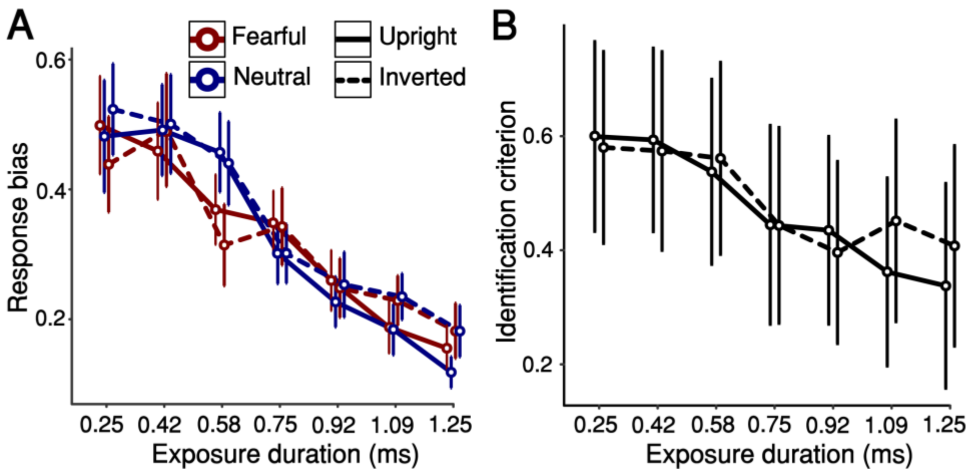


**Supplementary Fig. 11**. Additional results of Experiment 3. (A) Absolute-value response bias scores for reporting location (bias toward either left or right). The amount of response bias decreased as exposure duration increased. However, there was no statistically significant difference between fearful and neutral expressions or between upright and inverted faces. (B) Criterion scores for reporting expression. We did not find main effects or significant interactions. Data are presented as mean values with ±1 SEM bars, n = 32 independent participants.

#### Identification sensitivity

To examine how the manipulated factors affected expression identification, we entered identification d′ scores into a 2 (orientation: upright, inverted) × 7 (exposure durations) repeated-measures ANOVA. We did not find a main effect of exposure duration ($F_{(6, 186)} = 0.876, p = .513, \eta p^2 = .027$) or face orientation ($F_{(1, 31)} = 1.324, p = .259, \eta p^2 = .041$). Their interaction did not reach significance either ($F_{(4.31, 133.63)} = 0.486, p = .759, \eta p^2 = .015$). As shown in **Fig. 2B**, these results indicate that single-stimulus identification did not depart from chance with exposure durations ranging between 0.25 and 1.25 ms.

#### Expression identification criterion

We examined whether participants' criterion for reporting fearful expression varied across conditions by entering $C_{identification}$ scores into a 2 (orientation: upright, inverted) × 7 (exposure durations) repeated-measures ANOVA. We did not find a significant main effect of exposure duration ($F_{(1.59, 49.14)} = 2.824, p = .081, \eta p^2 = .083$), or a main effect of face orientation ($F_{(1, 31)} = 0.314, p = .579, \eta p^2 = .01$), or an interaction between them ($F_{(6, 186)} = 1.34, p = .241, \eta p^2 = .041$), (**Supplementary Fig. 11B**). These results suggest that identification criterion was unaffected by the experiment manipulations.

**Supplementary Note 8. Additional details of Experiment 4 and comparison of identification between Experiments 1 and 4**

**Location sensitivity**

To examine how the manipulated factors affected stimulation location detection, we entered location d′ scores into a 2 (expression: fearful, neutral) × 2 (orientation: upright, inverted) × 7 (exposure durations) repeated-measures ANOVA. We found a main effect of exposure duration ($F_{(1.276, 39.557)} = 143.034, p < .00001, \eta p^2 = .822$). As shown in **Fig. 2C**, location d′ scores increased to high sensitivity at the longest exposure durations, indicating that changes in exposure duration between 0.8 and 6.2 ms had great impact on participants' ability to detect stimulation, reaching ceiling in sensitivity with 1.7 ms. We did not find a main effect of face orientation ($F_{(1, 31)} = 1.452, p = .237, \eta p^2 = .045$) or emotional expression ($F_{(1, 31)} = 0.004, p = .953, \eta p^2 < 0.01$). No interaction reached significance either (all $p$ < .470).

**Location response bias**

We examined whether participants' response bias for reporting the face presentation location varied across conditions by entering the absolute values of $C_{order}$ scores into a 2 (expression: fearful, neutral) × 2 (orientation: upright, inverted) × 7 (exposure durations) repeated-measures ANOVA. Response bias significantly decreased with increasing exposure duration ($F_{(1.35, 41.76)} = 47.934, p < .00001, \eta p^2 = .607$), indicating that as participants' ability to detect stimulation increased, they became less likely to exhibit a systematic bias in their preference to report one presentation location or the other (**Supplementary Fig. 12A**). We did not find a main effect of expression ($F_{(1, 31)} = 0.349, p = .559, \eta p^2 = 0.01$), or of face orientation ($F_{(1, 31)} = 0.614, p = .439, \eta p^2 = .02$). The interaction between exposure duration and expression reached significance ($F_{(2.657, 82.365)} = 3.01, p = .041, \eta p^2 = 0.09$), but only the post-hoc pairwise comparison at 0.8 ms ($t(31) = 3.61, p = .035, d = .46, \mathrm{CI} = 0.002 - 0.166$) reached significance, with an advantage for fearful faces. No other interaction reached significance (all $p$ > .099). These results suggest that the magnitude of response bias decreased with exposure duration, with an advantage for fearful expressions at 0.8 ms of exposure.

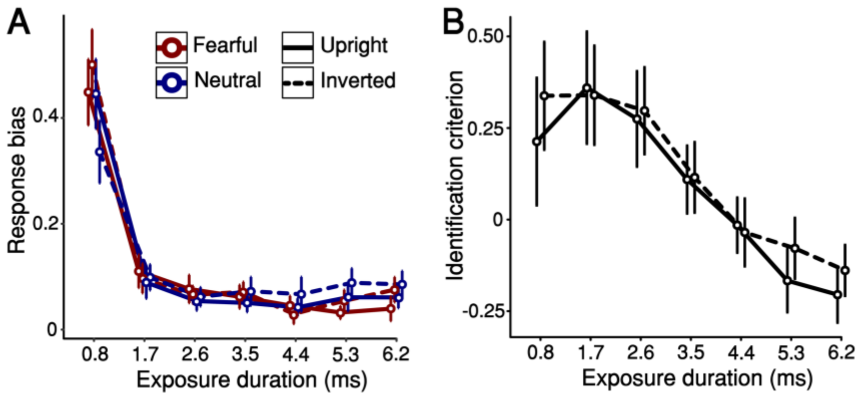


**Supplementary Fig. 12**. Additional results of Experiment 4. (A) Absolute-value response bias scores for reporting location (bias toward either left or right). The amount of response bias decreased as exposure duration increased, with slightly (but significantly) greater bias for inverted faces over upright faces. However, there was no statistically significant difference between fearful and neutral expressions (B) Criterion scores for reporting expression. Although we did not find main effects of exposure duration and face orientation, we did find a significant interaction between these two factors. However, post hoc comparisons did not reveal significant differences. Data are presented as mean values with ±1 SEM bars, n = 32 independent participants.

### Identification sensitivity

To examine how the manipulated factors affected expression identification, we entered identification d′ scores into a 2 (orientation: upright, inverted) × 7 (exposure durations) repeated-measures ANOVA. We found a main effect of exposure duration ($F_{(3.892,\ 120.665)} = 17.589, p < .00001, \eta p^2 = .362$), and a main effect of face orientation ($F_{(1,\ 31)} = 15.993, p < .00037, \eta p^2 = .340$). Their interaction also reached significance ($F_{(6,\ 186)} = 4.117, p = .00066, \eta p^2 = .117$). Bonferroni-corrected pairwise comparisons revealed a FIE arising at 5.3 ms ($t(31) = 4.014, p = .008, d = 0.972, \mathrm{CI} = 0.051 - 0.766$) and 6.2 ms of exposure ($t(31) = 5.162, p = .000053, d = 1.251, \mathrm{CI} = 0.168 - 0.883$). As shown in **Fig. 2D**, these results indicate that single-stimulus identification exhibits an advantage for upright faces over inverted faces from 5.3 ms of exposure.

### Expression identification criterion

We examined whether participants' criterion for reporting fearful expression varied across conditions by entering $C_{identification}$ scores into a 2 (orientation: upright, inverted) × 7 (exposure durations) repeated-measures ANOVA. We found a significant main effect of exposure duration ($F_{(2.05,\ 63.7)} = 9.804, p = .00017, \eta p^2 = .24$), indicating that as participants' ability to identify facial expression, their identification criterion became more liberal (**Supplementary Fig. 12B**). We did not find a main effect of face orientation ($F_{(1,\ 31)} = 0.295, p = .591, \eta p^2 = .009$) or an interaction between exposure duration and face orientation ($F_{(6,\ 186)} = 0.869, p = .519, \eta p^2 = .027$). These results suggest that identification criterion became more liberal as identification sensitivity increased.

### Comparison between Experiments 1 and 4 in identification sensitivity

To test whether the presence or absence of scrambled stimuli had any impact on expression identification, we compared identification sensitivity in Experiments 1 and 4 with a mixed ANOVA, treating the experiment as between-subject factor, and exposure duration and face orientation as within-subject factors. We found a main effect of exposure duration ($F_{(4.18, 251)} = 30.12, p < .00001, \eta p^2 = .327$), and a main effect of face orientation ($F_{(1, 62)} = 33.94, p < .00001, \eta p^2 = .354$). We also found a significant interaction between exposure duration and face orientation ($F_{(6, 372)} = 6.61, p < .00001, \eta p^2 = .096$). Crucially, we did not find an effect of Experiment ($F_{(1, 62)} = 0.012, p = .912, \eta p^2 < 0.01$). No other interaction reached significance (all $p > .497$). Post hoc pairwise comparisons revealed a significant advantage of upright faces over inverted faces at 5.3 ($t(63) = 5.349, p < .00001, d = 0.871, \mathrm{CI} = 0.119 - 0.562$) and 6.2 ms of exposure ($t(63) = 6.983, p < .00001, d = 1.138, \mathrm{CI} = 0.223 - 0.67$). See **Supplementary Fig. 13** for the identification sensitivity results of both experiments superimposed on the same axes. These results indicate that the presence or absence of scrambles did not influence expression identification sensitivity.

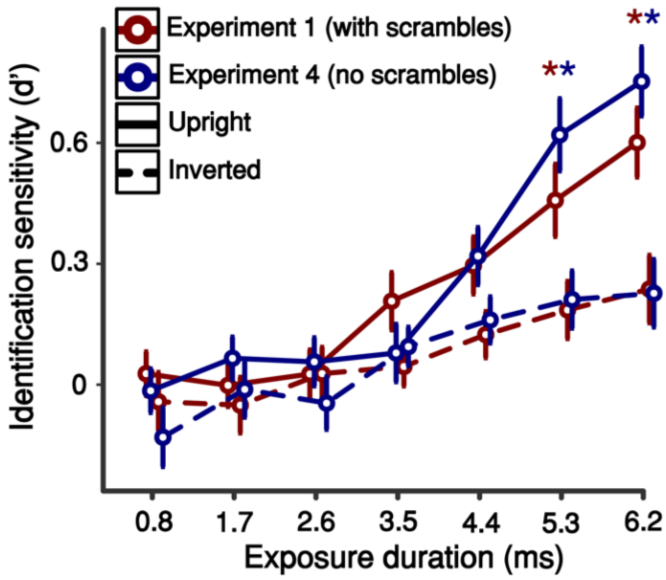


**Supplementary Fig. 13.** Results from Experiment 1 and Experiment 4's identification sensitivity. Experiment 1's expression identification results: Expression identification increased with exposure duration, and an advantage for upright faces was found from 5.3 ms of exposure. Experiment 4's expression identification results: The same pattern of results as in Experiment 1 was found. We found no differences between Experiment 1 and Experiment 4, which indicates that the presence of scrambles did not play a role in expression identification. Upright faces are indicated by solid lines, and inverted faces are indicated by dashed lines. Data are presented as mean values with ±1 SEM bars, n = 32 independent participants per experiment. Results from Experiment 1 are represented by red lines, while results from Experiment 4 are represented by blue lines. * $p < 0.05$ for upright-inverted comparisons, where comparisons for Experiment 1 are shown in red, and for Experiment 4 in blue.

### Supplementary Note 9: PAS ratings in Experiments 3 and 4

The PAS ratings obtained in Experiments 3 and 4 precluded the calculation of metacognitive sensitivity. Calculating meta-d′ requires a diverse range of PAS ratings for maximum-likelihood estimate model-fitting, with a sufficiently large number of correct and incorrect trials for each rating[8,40]. Participants provided too few high (almost clear / clear experience) ratings for all exposure durations in Experiment 3 (**Supplementary Fig. 14A**), and for most durations in Experiment 4 (**Supplementary Fig. 14B**). PAS rating distributions thus lacked an adequate spread for fitting the model-based estimates of metacognitive sensitivity. Only 6.3% (Experiment 3) and 21.9% (Experiment 4) of the participants′ data could be fitted for all conditions. This contrasts with the PAS ratings obtained in Experiments 1 and 2, wherein all participants′ data could be fitted to the MLE meta-d′ model for all conditions.

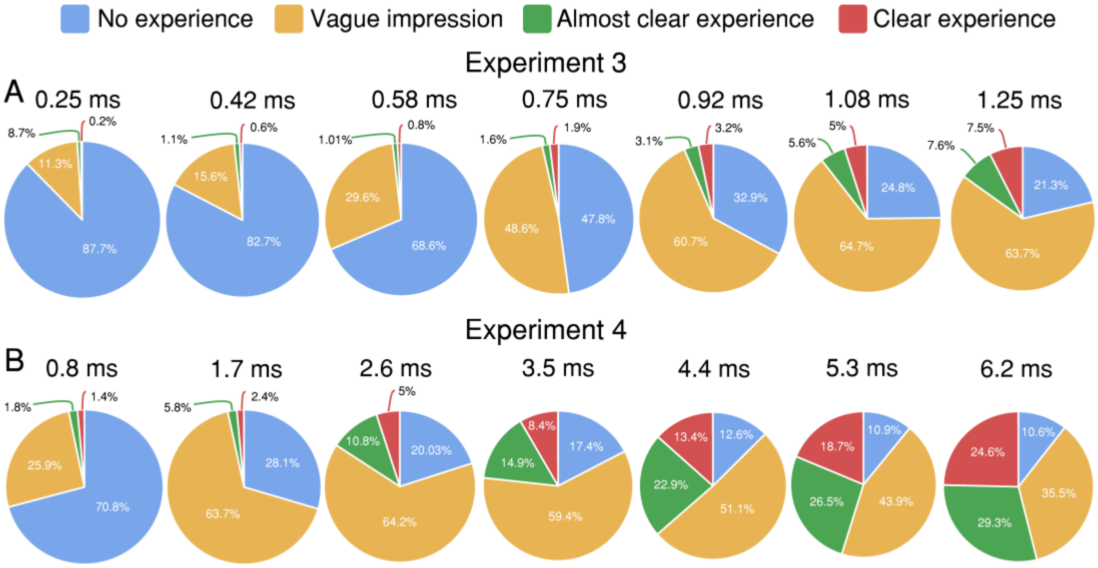


**Supplementary Fig. 14.** Mean PAS rating frequencies for Experiments 3 and 4. (A) In Experiment 3, most exposure durations yielded almost no ‘almost clear experience’ and ‘clear experience’ ratings. (B) In Experiment 4, most exposure durations resulted in very low 'almost clear experience' and 'clear experience' ratings across several durations. Percentages represent overall numbers per exposure durations.

### Supplementary Note 10: Power analysis, face stimuli, and EEG analysis in Experiment 5

#### Participants

Thirty-six Université Libre de Bruxelles students provided informed consent and were paid €30 for participation. All had normal or corrected-to-normal vision and reported no history of neurological or psychiatric disorders. Four participants were excluded from the analysis (see Analysis section). The remaining 32 participants (18 female; all right-handed) had a mean age of 24.3 ($SD_{age}$ = 4.9; range: 18 – 31).

An a priori power analysis, conducted using G*Power 3.1.9.7[41], to test for a difference between conditions (main effects of orientation and expression) in a repeated-measures ANOVA, with a small to medium effect size ($\eta p^2 = 0.04$) and alpha of .05, aiming to achieve a statistical power of 95%, determined that a sample of 19 participants would be required. If a non-sphericity correction ε of .5 were to be added – as reported in the Results section, a number of tests violated this assumption – then a sample of 29 participants would be required. This analysis supports our initial decision of aiming at 32 participants per experiment.

#### Stimuli

Stimuli were 60 human faces (20 fearful, 20, happy, and 20 neutral; the same 20 identities – 10 female – were used for the three categories; **Supplementary Fig. 15**) taken from the RaFD. They were selected by applying the same criteria used in Experiment 1; their differences in expression identification ($M_{\text{fearful}}$ = 92.2%; [$SD_{\text{fearful}}$ = 5.33]; $M_{\text{happy}}$ = 99.15% [1.76]; $M_{\text{neutral}}$ = 95.15% [5.73]) and intensity ($M_{\text{fearful}}$ = 4.22; [0.21]; $M_{\text{happy}}$ = 4.20 [0.26]; $M_{\text{neutral}}$ = 3.61 [0.25]) were minimised. The same image processing steps were followed, too.

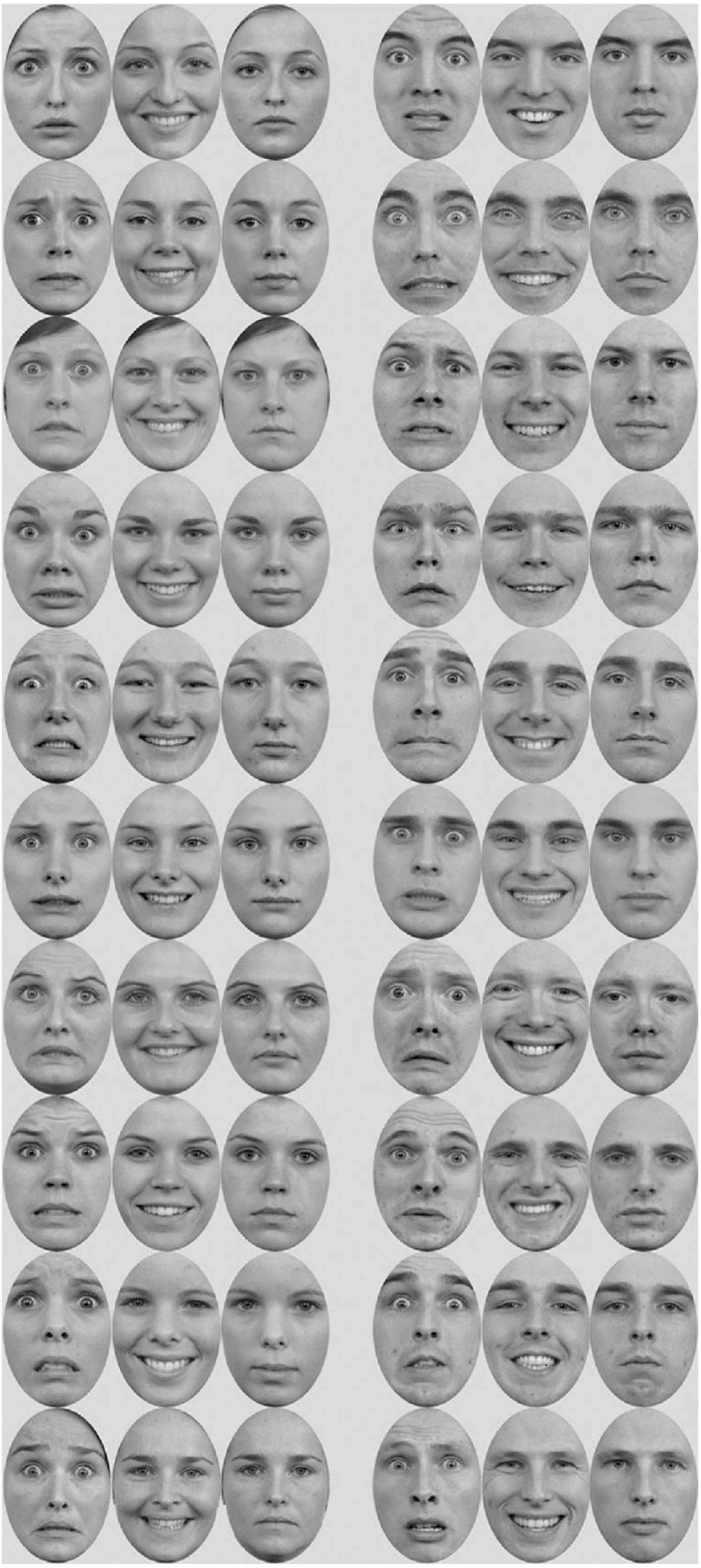

**Supplementary Fig. 15.** RaFD stimuli used in Experiment 5: fearful, happy, and neutral faces. Female faces on the left, male faces on the right. The intact faces shown here belong to the Radboud Face Database (RaFD) and can be presented as stimulus examples (see: https://rafd.socsci.ru.nl/).

A one-way ANOVA was conducted to compare expression identification between emotional expressions (fearful, happy, neutral) based on the norms published by Langner et al.[3]. The effect of emotional valence was significant ($F_{(2, 57)} = 11.13, p = .00007, \eta^2p = .285$). Bonferroni-corrected pairwise comparisons revealed that happy expressions were easier to identify than fearful ($t(59) = 4.746, p = .000043, d = 1.5, \mathrm{CI} = -10.562 - -3.338$) and neutral expressions ($t(59) = 2.732, p = .025, d = 0.864, \mathrm{CI} = 0.388 - 7.612$). However, we did not find a significant difference in identification between fearful and neutral expressions ($t(59) = 2.95, p = .146, d = 0.637, \mathrm{CI} = -6.562 - 0.662$). See **Supplementary Fig. 16A**.

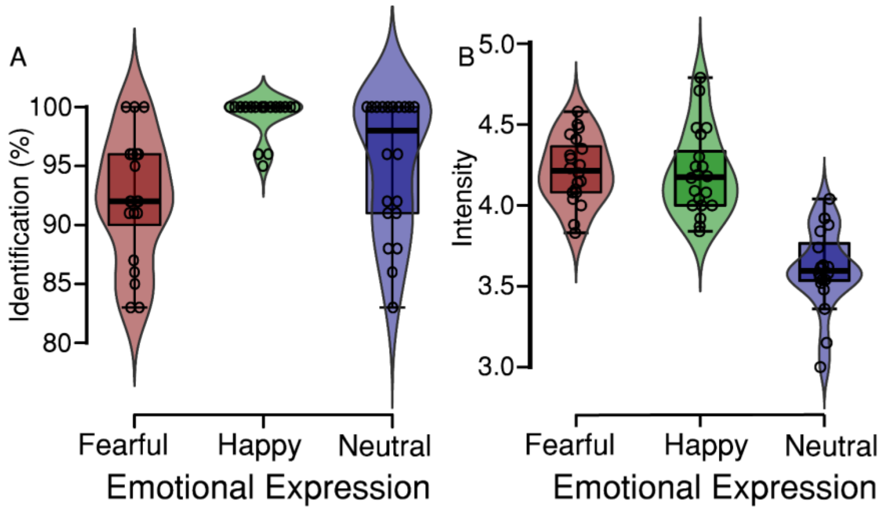


**Supplementary Fig. 16.** (A) Expression identification/agreement (%) scores and (B) Expression identity scores of RaFD stimuli selected for Experiment 5, based on RaFD validation norms, between emotional expressions (fearful, happy, neutral). Boxplots show median, interquartile range, and density curves depicting data distribution. Individual dots represent individual ratings.

A one-way ANOVA was conducted to compare emotional intensity between emotional expressions (fearful, happy, neutral) based on the same norms. The effect of emotional valence was significant ($F_{(2, 57)} = 41.56, p < .00001, \eta^2p = .593$). Bonferroni-corrected pairwise comparisons revealed that fearful expressions were rated as more intense than neutral expressions ($t(59) = 7.985, p < .00001, d = 2.525, \mathrm{CI} = 0.422 - 0.8$), but not than happy expressions ($t(59) = 0.183, p = 1, d = 0.058, \mathrm{CI} = -0.175 - 0.203$). Likewise, happy expressions were rated as more intense than neutral expressions ($t(59) = 7.802, p < .00001, d = 2.467, \mathrm{CI} = 0.408 - 0.786$). See **Supplementary Fig. 16B**.

**EEG analysis**

Experiments 1 (2AFC) and 2 (2IFC) had found a face-inversion effect (FIE, a perceptual index of holistic face processing) on location d'. However, the psychophysical measures of those behavioural experiments did not reveal any effects of emotion on detection. The aim of Experiment 5 was therefore to search for *neural* markers of emotion processing. To this end, we included three emotional expressions with different valences, to enable multiclass decoding analysis (see **Fig. 4J-K**). In light of the large number of trials required for this analysis, which necessitated a long experiment (3-4 hours), we did not include inverted faces in Experiment 5. Unlike the neural indices of emotion processing, obtaining neural measures of the FIE would have added little to the FIE results we already had from Experiments 1 and 2.

The event-related potential (ERP) analysis was implemented to determine the minimal duration required for neural markers of emotional processing of facial expressions to arise. We further examined the effect of increasing exposure durations on these markers. The selection of ERP components was based on a well-established literature comprising a very large number of studies that have validated the sensitivity of these measures to the stimulus attributes of interest and described both their typical scalp topography and latency. The P1 component is sensitive to low-level visual information and visual attention, and is typically measured 100 ms after stimulus onset around occipital electrodes[42]. The N170/VPP component is sensitive to face stimuli compared to non-face stimuli (most typically, object images), and it is typically found around occipital electrodes between 170 and 200 ms after stimulus onset[43–49]; its source lies in the fusiform gyrus (roughly, around the fusiform face area[50]). The early posterior negativity (EPN) component is sensitive to differences in early attentional capture between emotional and neutral images, and it is typically found over occipital electrodes between 150 and 300 ms after stimulus onset[51–58]. The late-positive potential (LPP) component is sensitive to differences in emotional intensity between visual stimuli, and it is typically found over occipital areas between 300 and 400 ms after stimulus onset[52,59–68]. The Visual awareness negativity (VAN), also known as the perceptual awareness negativity[69], is sensitive to awareness and is typically found around occipital electrodes around 200 ms after stimulus onset[70–73]. Late positivity (LP) is typically found over centro-parietal areas between 300 and 500 ms after stimulus onset[71,74]. We did not analyse evoked EEG data outside of canonical electrodes and latencies for each ERP component of interest.

### Supplementary Note 11: Full Results of Experiment 5

#### Behavioural Results

#### Location sensitivity

Neutral expressions appeared in both fearful and happy expression blocks. We entered location d′ scores for neutral-expression trials into a preliminary repeated-measures 2 (Block-type: neutral-expression in fearful vs happy expression blocks) × 3 (exposure durations) repeated-measures ANOVA; there was no significant effect of block type ($F_{(1, 31)} = 0.358, p = .554, \eta p^2 = .011$), nor an interaction of block type with exposure duration ($F_{(1.945, 60.291)} = 1.468, p = .239, \eta p^2 = .045$). We also estimated Bayes factors: the results suggest the data are substantially better explained under the null hypothesis model ($BF_{01} = 6.440$). We therefore collapsed neutral-expression trials into one condition, making three conditions in total.

In the main analysis, we measured location d′ to confirm that these new participants displayed similar performance to participants in Experiment 1. To examine how conditions affected location discrimination, we entered location d′ scores into a 3 (expression: fearful, happy, neutral) × 3 (exposure durations) repeated-measures ANOVA. We found a main effect of exposure duration, whereby sensitivity increased with increasing durations ($F_{(2, 62)} = 345.823, p < .00001, \eta p^2 = .918$). Importantly, we found very similar location d′ scores to the ones reported in Experiment 1 (**Supplementary Fig. 17A**). As expected, based on the findings of Experiment 1, we did not find a main effect of expression ($F_{(2, 62)} = 0.397, p = .674, \eta p^2 = .013$), thus suggesting again that there is no greater perceptual sensitivity to emotional expressions than to neutral ones. The interaction between expression and exposure duration did not reach significance either ($F_{(4, 124)} = 1.711, p = .152, \eta p^2 = .052$). Thus, location sensitivity performance replicated the findings obtained in equivalent conditions in Experiment 1.

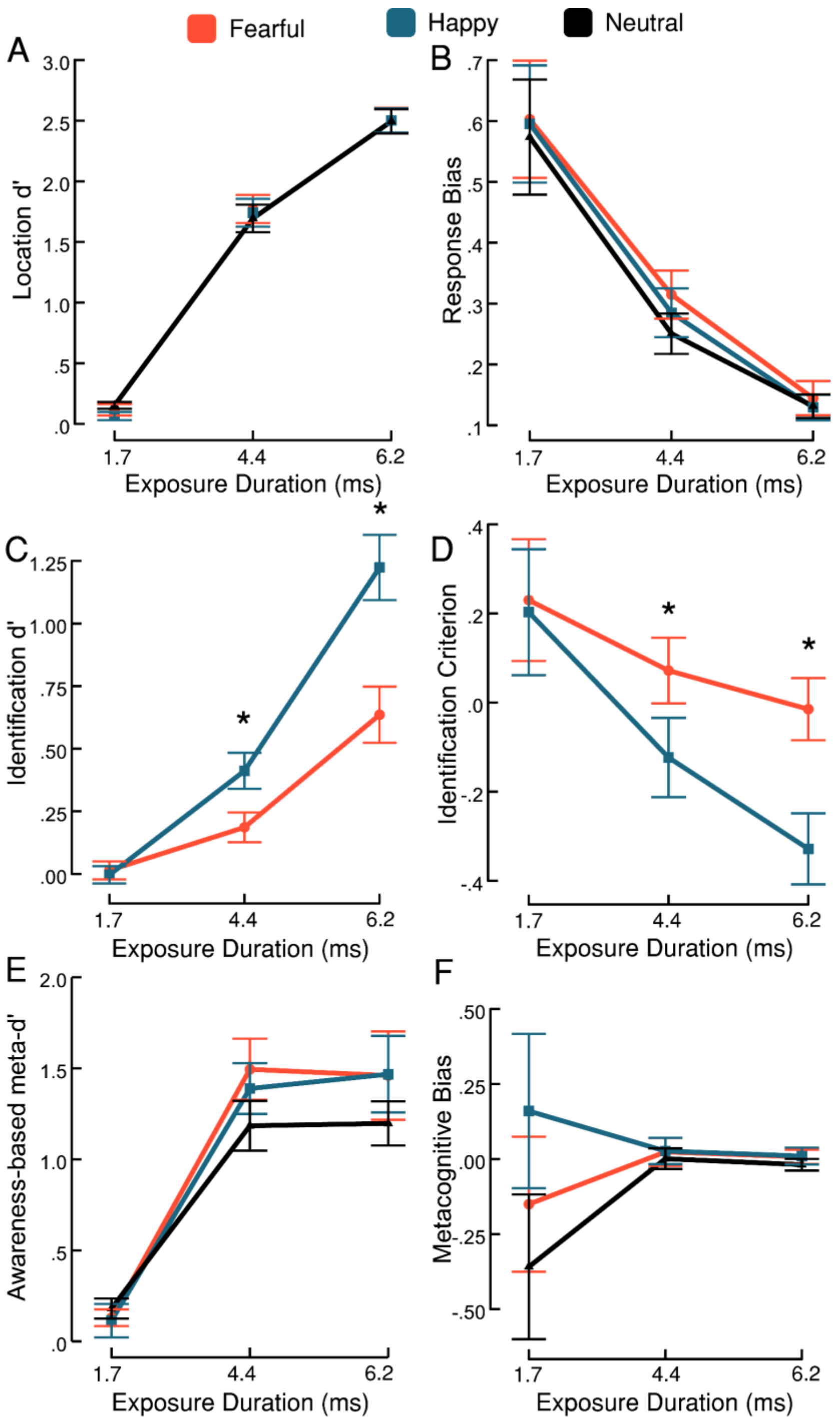


**Supplementary Fig. 17.** Full behavioural results of Experiment 3. (A) Location sensitivity. A repeated-measures ANOVA showed that location d′ increased with increasing exposure duration. There was no effect of emotional expression. (B) Absolute-value response bias scores for reporting location (bias toward either left or right). A repeated-measures ANOVA showed that the amount of bias decreased as exposure duration increased, but there was no difference in amount of response bias between expressions. (C) Identification sensitivity for expression. A repeated-measures ANOVA showed that identification d′ increased with increasing exposure duration. A significant advantage in expression identification for happy expressions over fearful expressions arises by 4.4 ms of exposure. (D) Criterion scores for reporting expression. A repeated-measures ANOVA showed that criterion becomes more liberal with increasing exposure duration. A more liberal criterion for happy expressions over fearful expressions arises by 4.4 ms of exposure. (E) Awareness-based metacognitive sensitivity. A repeated-measures ANOVA showed that meta-d′ increased with exposure duration but was unaffected by expression. (F) Metacognitive bias scores for reporting subjective awareness. A repeated-measures ANOVA showed that metacognitive bias was unaffected by exposure duration and expression. * $p < 0.05$ for fearful-happy comparisons. Data are presented as mean values with ±1 SEM bars, n = 32 independent participants.

To determine the shortest exposure that exhibited above-chance performance (d′ > 0), we ran a series of uncorrected one-sample t-tests against zero. We found that above-chance identification was already present at the shortest exposure duration, 1.7 ms, for fearful expressions ($M = 0.120$ [0.285]; $t(31) = 2.47, p = .019, d = 0.436, \mathrm{CI} = 0.02 - 0.212$), neutral expressions ($M = 0.172$ [0.208]; $t(31) = 5.41, p = .000066, d = 0.956, \mathrm{CI} = 0.095 - 0.21$), and happy expressions ($M = 0.070$ [0.204]; $t(31) = 1.922, p = .06, d = 0.34, \mathrm{CI} = -0.004 - 0.132$). Thus, above-chance performance was found at slightly shorter exposure durations than in Experiment 1.

#### Location response bias

We examined whether participants′ response bias for reporting face location varied across conditions by entering the absolute values of $C_{location}$ scores into a 3 (emotional expression: fearful, happy, neutral) × 3 (exposure durations) repeated-measures ANOVA. As in Experiment 1, response bias significantly decreased with increasing exposure duration ($F_{(1.09, 33.78)} = 20.1, p < .00001, \eta p^2 = .393$), indicating that as participants′ ability to discriminate faces increased, they became less likely to exhibit a systematic bias in their preference to report one side or the other (**Supplementary Fig. 17B**). However, we did not find a main effect of expression ($F_{(2, 62)} = 2.061, p = .136, \eta p^2 = .062$), suggesting that response bias was unaffected by the emotional content of the faces. The interaction between expression and exposure duration did not reach significance either ($F_{(4, 124)} = 0.571, p = .684, \eta p^2 = .018$).

To assess whether the obtained data support the absence of an effect of emotional expression, we estimated the Bayes factor for this effect, which indicated strong evidence for the null hypothesis model ($BF_{01} = 16.978$).

#### Expression identification sensitivity

We examined whether participants′ sensitivity to identifying emotional expressions from their neutral counterparts varied across conditions by entering identification d′ scores into a 2 (emotional expression: fearful, happy) × 3 (exposure durations) repeated-measures ANOVA (**Supplementary Fig. 17C**). As expected, we found a main effect of exposure duration ($F_{(1.38, 42.74)} = 62.927, p < .00001, \eta p^2 = .670$), indicating that sensitivity to emotional expression increased with increasing exposure duration. We also found a main effect of expression ($F_{(1, 31)} = 36.25, p = .0000012, \eta p^2 = .539$), showing better sensitivity to happy expressions ($M = 0.544$ [0.625]) than to fearful expressions ($M = 0.278$ [0.321]). The interaction between expression and exposure duration also reached significance ($F_{(1.65, 51.01)} = 25.569, p < .00001, \eta p^2 = .452$). To determine at which of the three durations we used this advantage was present, we ran post hoc Bonferroni-corrected pairwise comparisons. They revealed significant advantages at 4.4 ms ($t(31) = 3.416, p = .014, d = 0.485, \mathrm{CI} = -0.425 - -0.026$) and 6.2 ms of exposure ($t(31) = 8.901, p < .00001, d = 1.262$).

To determine the shortest exposure that exhibited above-chance performance (d′ > 0), we ran a series of uncorrected one-sample t-tests against zero. We found that the earliest exposure duration that elicited above-chance identification was 4.4 ms for fearful ($M = 0.186$ [0.338]; $t(31) = 3.113, p = .004, d = 0.550, \mathrm{CI} = 0.064 - 0.308$) and happy facial expressions ($M = 0.412$ [0.406]; $t(31) = 5.733, p = .0000026, d = 1.014, \mathrm{CI} = 0.265 - 0.558,$). Neither expression exhibited above-chance performance with 1.7 ms of exposure, therefore we estimated Bayes factors to test whether the obtained data support this null effect. Bayes factors indicated substantial evidence in favour of the null hypothesis model for fearful ($BF_{01} = 4.93$) and happy expressions ($BF_{01} = 5.27$) presented for 1.7 ms of exposure, thus supporting the finding that neither expression enjoyed above-chance performance when presented for 1.7 ms.

These findings, along with the metacognitive sensitivity findings reported below, will be relevant for our EEG analysis – if we were to find neural evidence of emotion processing with 1.7 ms of exposure by measuring neural markers, this would suggest unconscious emotion processing.

**Expression identification criterion**

We examined whether participants′ criterion for reporting fearful and happy expressions varied across conditions by entering $C_{identification}$ scores into a 2 (expression: fearful, happy) × 3 (exposure durations) repeated-measures ANOVA. We found a main effect of exposure duration ($F_{(1.11, 34.32)} = 4.664, p = .034, \eta p^2 = .131$), indicating that identification criterion became more liberal across increasing exposure duration, i.e., participants became more willing to report an emotional expression (fearful or happy) than a neutral one as exposure duration increased (**Supplementary Fig. 17D**). We also found a main effect of emotional expression ($F_{(1, 31)} = 15.418, p = .00045, \eta p^2 = .332$), indicating that happy expressions ($M = -0.097$ [0.226]) enjoyed a more liberal identification criterion than fearful expressions ($M = 0.079$ [0.086]). Finally, the interaction between both factors reached significance as well ($F_{(1.62, 50.228)} = 17.567, p = .0000071, \eta p^2 = .362$). To determine at which exposure durations happy expressions enjoyed a significantly more liberal criterion than fearful expressions, we ran post hoc Bonferroni-corrected pairwise comparisons. They revealed significant advantages at 4.4 ms ($t(31) = 3.635, p = .009, d = 0.328, CI = 0.03 - 0.352$) and 6.2 ms of exposure ($t(31) = 5.874, p = .0000039, d = 0.530, \mathrm{CI} = 0.147 - 0.470$).

**Awareness-based metacognitive sensitivity**

We examined whether awareness scores were sensitive to participants′ location sensitivity scores by estimating meta-d′, a measure of metacognitive sensitivity. To examine whether meta-d′ varied across conditions, we entered meta-d′ scores into a 3 (expression: fearful, happy, neutral) × 3 (exposure durations) repeated-measures ANOVA (**Supplementary Fig. 17E**). A main effect of exposure duration indicated that meta-d′ increased with increasing exposure duration ($F_{(2, 62)} = 55.970, p < .00001, \eta p^2 = .644$). However, we did not find a main effect of expression ($F_{(2, 62)} = 1.297, p = .281, \eta p^2 = .040$), suggesting that emotional expression did not affect metacognitive sensitivity. To examine the duration interval in which awareness arose, we ran post hoc Bonferroni-corrected pairwise comparisons between each combination of exposure durations (collapsed across expressions). We found that meta-d′ scores obtained with 4.4 ms were significantly higher than scores obtained with 1.7 ms ($t(31) = 9.09, p < .00001, d = 1.449, \mathrm{CI} = -1.543 - -0.886$), whereas meta-d′ scores obtained with 6.2 ms were not significantly higher than scores obtained with 4.4 ms ($t(31) = 0.143, p = .999, d = 0.023, \mathrm{CI} = -0.348 - 0.310$). This may suggest that 4.4 ms of exposure provided sufficient visual information for meta-d′ to exhibit a non-linear increase. Finally, we did not find an interaction between expression and exposure duration ($F_{(2.69, 83.32)} = 0.934, p = .420, \eta p^2 = .029$).

As described, we did not find a main effect of expression, therefore we calculated Bayes factors to test whether the obtained data support this absence of an effect. Bayes factors indicated strong evidence in favour of the null hypothesis model ($BF_{01} = 12.06$), thus supporting the finding that no expression was prioritised by metacognitive sensitivity.

To determine the shortest exposure that exhibited above-chance metacognitive sensitivity (meta-d′ > 0), we ran a series of uncorrected one-sample t-tests against zero. We found that above-chance meta-d′ was already present at the shortest exposure duration, 1.7 ms, for fearful ($M = 0.130$ [0.259]; $t(31) = 2.843, p = .008, d = 0.503, \mathrm{CI} = 0.037 - 0.223$) and neutral expressions ($M = 0.181$ [0.313]; $t(31) = 3.268, p = .003, d = 0.578$); meta-d′ was above chance at 4.4 ms for happy expressions ($M = 1.389$ [0.790]; $t(31) = 9.951, p < .00001, d = 1.759, \mathrm{CI} = 1.104 - 1.674$). Although meta-d′ scores were higher than zero at the shortest exposure duration used, they were not higher than location d′ scores at the same exposure duration.

#### Metacognitive bias

Meta-bias is the tendency to give high confidence ratings regardless of actual performance. Participants described their visual experience of a face and, therefore, meta-bias here describes the tendency to describe one's visual experience as clear. To examine whether meta-bias varied across conditions, we entered meta-bias scores into a 3 (expression: fearful, happy, neutral) × 3 (exposure durations) repeated-measures ANOVA (**Supplementary Fig. 17F**). We did not find an effect of exposure duration ($F_{(1.05, 32.51)} = 0.555, p = .470, \eta p^2 = .018$), or of facial expression ($F_{(2, 62)} = 1.585, p = .213, \eta p^2 = .049$). The interaction between expression and exposure duration did not reach significance either ($F_{(1.97, 61.17)} = 1.226, p = .30, \eta p^2 = .038$). These results suggest that the participants' tendency to report a clear visual experience was not affected by any of the conditions.

### EEG results

### P1

We examined how emotional expression and exposure duration conditions affected early visual processing by measuring voltage changes in P1. This component is measured over occipital regions. We entered mean voltage values into a 2 (expression: emotional, neutral) ×

3 (exposure durations) repeated-measures ANOVA. We found a main effect of exposure duration ($F_{(1.14, 35.27)} = 154.18, p < .00001, \eta p^2 = .833$), whereby P1 voltage increased with exposure duration, thus indicating that longer exposure durations involved greater early visual processing than shorter exposure durations (**Supplementary Fig. 18**). As expected, given the nature of this component, we did not find a main effect of expression ($F_{(1, 31)} = 1.21, p = .280, \eta p^2 = .038$) and the interaction between expression and exposure duration did not reach significance either ($F_{(2, 62)} = 2.026, p = .140, \eta p^2 = .061$). To assess whether the obtained data support the absence of an effect of emotional expression, we estimated a Bayes factor, which indicated substantial evidence for the null hypothesis model ($BF_{01} = 6.385$). These results indicate that P1 is sensitive to extremely brief visual exposure durations but is not sensitive to emotional expressions.

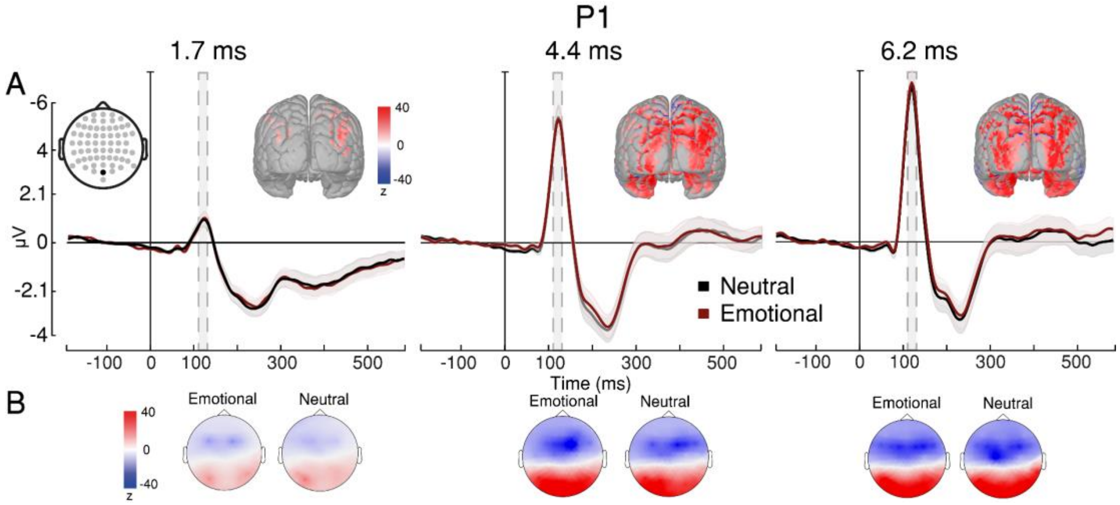


**Supplementary Fig. 18.** Early visual processing indexed by P1. (A) P1 evoked response to emotional expressions is represented by red lines, while evoked response to neutral expressions is represented by black lines; results shown for 1.7 ms (left), 4.4 ms (centre), and 6.2 ms of exposure (right). P1 peak with averaged time window highlighted in grey (105–135 ms) and source estimation of P1 visually identified on cortical maps (z-scale: from -40 to 40). (B) Topographical distributions of P1 for each condition at the relevant time window show an increase in positive voltage over visual cortex across exposure durations (z-scale: from -40 to 40). Shaded contours represent ±1 SEM, n = 32 independent participants. * $p < 0.05$ for emotional-neutral comparisons.

### N170/VPP

We examined how facial expression and exposure duration conditions affected face processing by measuring voltage changes in the N170/VPP complex. N170/VPP consists of two negative peaks (left and right N170) found in occipitotemporal areas, bilaterally, and one positive peak (vertex positive potential or VPP) found in frontocentral areas. We entered mean voltage values into a 2 (expression: emotional, neutral) × 3 (electrode site: left, right, central) × 3 (exposure durations) repeated-measures ANOVA. We found a main effect of exposure duration ($F_{(1.10, 34.20)} = 43.195, p < .00001, \eta p^2 = .582$), whereby N170/VPP amplitude changed with increasing exposure duration, thus suggesting longer exposure durations involved greater face processing than shorter exposure durations (**Supplementary Fig. 19**). We also found a main effect of electrode site ($F_{(1.49, 46.24)} = 94.125, p < .00001, \eta p^2 = .752$), which was expected given that left ($M = -5.404$ [1.674]) and right ($M = -5.554$ [1.523]) N170 potentials are negative-voltage peaks whereas VPP is a positive-voltage peak ($M = 4.575$ [1.085]). This effect confirmed that we measured the N170/VPP complex. We also found a main effect of expression ($F_{(1, 31)} = 11.696, p = .002$, $\eta p^2 = .274$), indicating that neutral expressions ($M = -2.198$ [5.303]) had significantly more negative voltage values than emotional expressions ($M = -2.058$ [5.134]). However, the interaction between expression and exposure duration did not reach significance ($F_{(1.57, 48.65)} = 0.318, p = .676$, $\eta p^2 = .01$). In addition, we found a significant interaction between expression and electrode site ($F_{(1.37, 42.32)} = 4.716, p = .025$, $\eta p^2 = .132$), which was expected given electrode sites' opposite voltage polarities, and a significant interaction between electrode site and exposure duration ($F_{(1.74, 53.95)} = 56.074, p < .00001$, $\eta p^2 = .644$), also expected given that N170 sites and VPP site became more negative and positive, respectively, as exposure duration increased. The three-way interaction between expression, electrode site, and exposure duration did not reach significance. Together, these results indicate that both exposure duration and expression affect N170/VPP – greater duration increases its amplitude (more negative for N170 and more positive for VPP), whereas the presence of emotion decreases it (less negative for N170 and less positive for VPP).

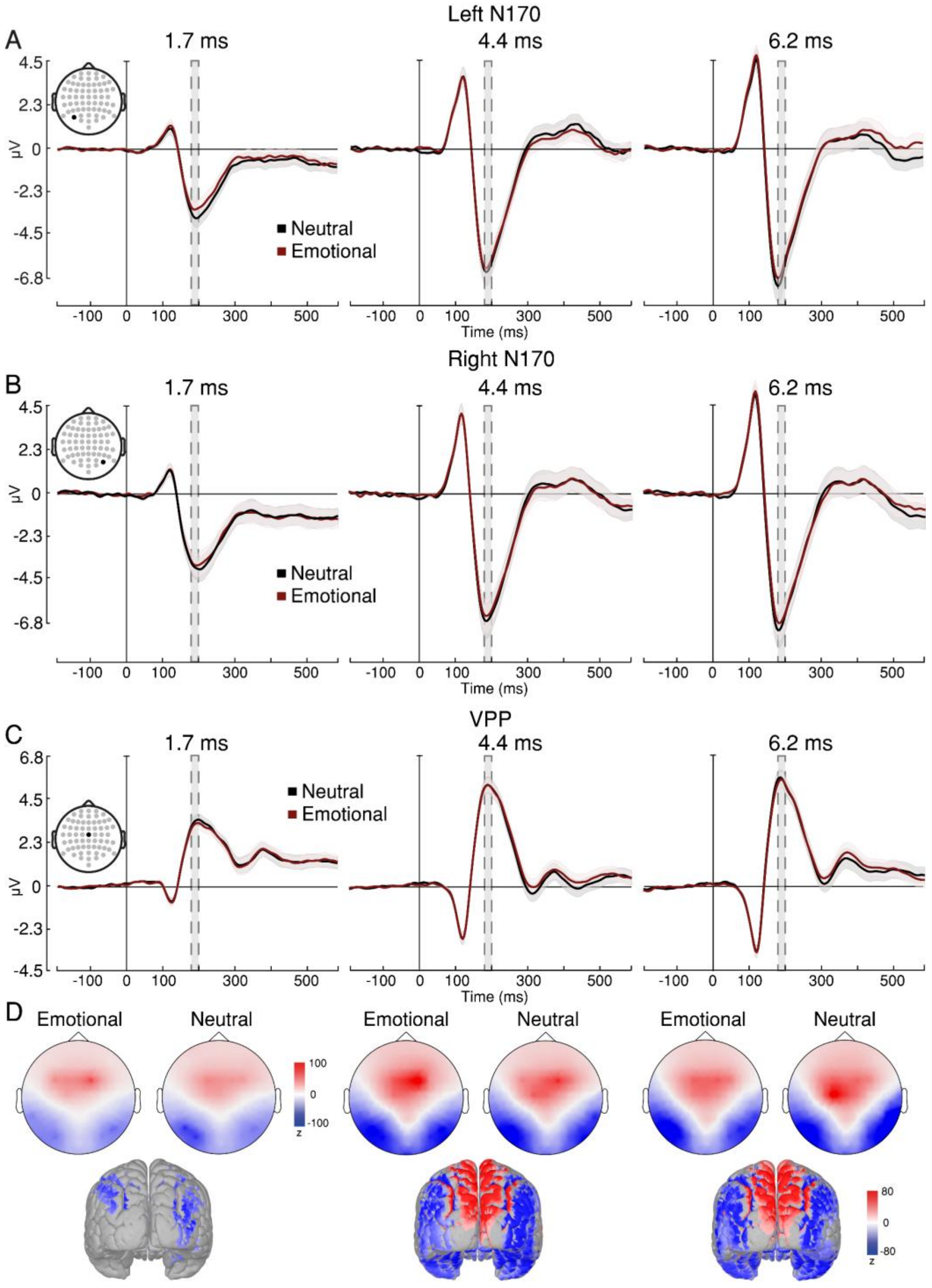


**Supplementary Fig. 19.** Face processing indexed by N170/VPP. (A) Left N170, (B) right N170, and (C) VPP response to emotional and neutral expressions across exposure durations. Evoked response to emotional expressions is represented by red lines, while evoked response to neutral expressions is represented by black lines. N170/VPP peak with averaged time window highlighted in grey (170–200 ms). (d) Topographical distributions of N170/VPP for each condition at the relevant time window (z-scale: from -100 to 100). Source estimation of N170/VPP visually identified on cortical maps (z-scale: from -80 to 80). Shaded contours represent ±1 SEM, n = 32 independent participants.

**Early Posterior Negativity (EPN)**

We examined how emotional expression and exposure duration conditions affected early emotion processing by measuring voltage changes in the early posterior negativity (EPN) component. This component is measured over occipitotemporal regions, bilaterally. We entered mean voltage values into a 2 (expression: emotional, neutral) × 2 (electrode site: left, right) × 3 (exposure durations) repeated-measures ANOVA. We did not find an effect of exposure duration ($F_{(1.26, 39.18)} = 2.884, p = .089, \eta p^2 = .085$), indicating that EPN did not vary across exposure durations overall (**Supplementary Fig. 20**). Importantly, however, we found a main effect of expression ($F_{(1, 31)} = 6.042, p = .02, \eta p^2 = .163$), thus confirming EPN was sensitive to emotional content in faces, showing more negative values for emotional ($M = -2.166$ [0.374]) than neutral faces ($M = -1.981$ [0.444]). We did not find an effect of electrode site ($F_{(1, 31)} = 2.157, p = .152, \eta p^2 = .065$), indicating EPN responded to changes across conditions similarly in both hemispheres. Crucially, we found a significant interaction between emotional expression and exposure duration ($F_{(2, 62)} = 8.675, p = .00048, \eta p^2 = .219$), suggesting EPN was sensitive to emotional content at specific exposure durations. To test at what specific exposure durations EPN was sensitive to emotional content, we ran post hoc Bonferroni-corrected pairwise comparisons and found that EPN was significantly more negative for emotional expressions ($M_{6.2\text{ms}} = -2.793$ [2.580]) than neutral expressions ($M_{6.2\text{ms}} = -2.070$ [3.423]) only at the longest exposure duration ($t(31) = 4.009, p = .002, d = 0.162, \text{CI} = -0.791 - -0.112$), when both awareness and emotion identification are very likely; therefore, these results suggest that emotion processing requires a longer exposure duration than holistic face processing to occur. Less importantly, we found a significant interaction between electrode site and exposure duration ($F_{(1.49, 46.14)} = 5.441, p = .013, \eta p^2 = .149$), though no comparison of interest (i.e., between electrode sites for each exposure duration) reached significance. The three-way interaction between expression, electrode site, and exposure duration did not reach significance either ($F_{(2, 62)} = 0.038, p = .963, \eta p^2 = .001$). These results suggest that emotion processing arises with exposure durations that are greater than 4.4 ms but smaller than or equal to 6.2 ms.

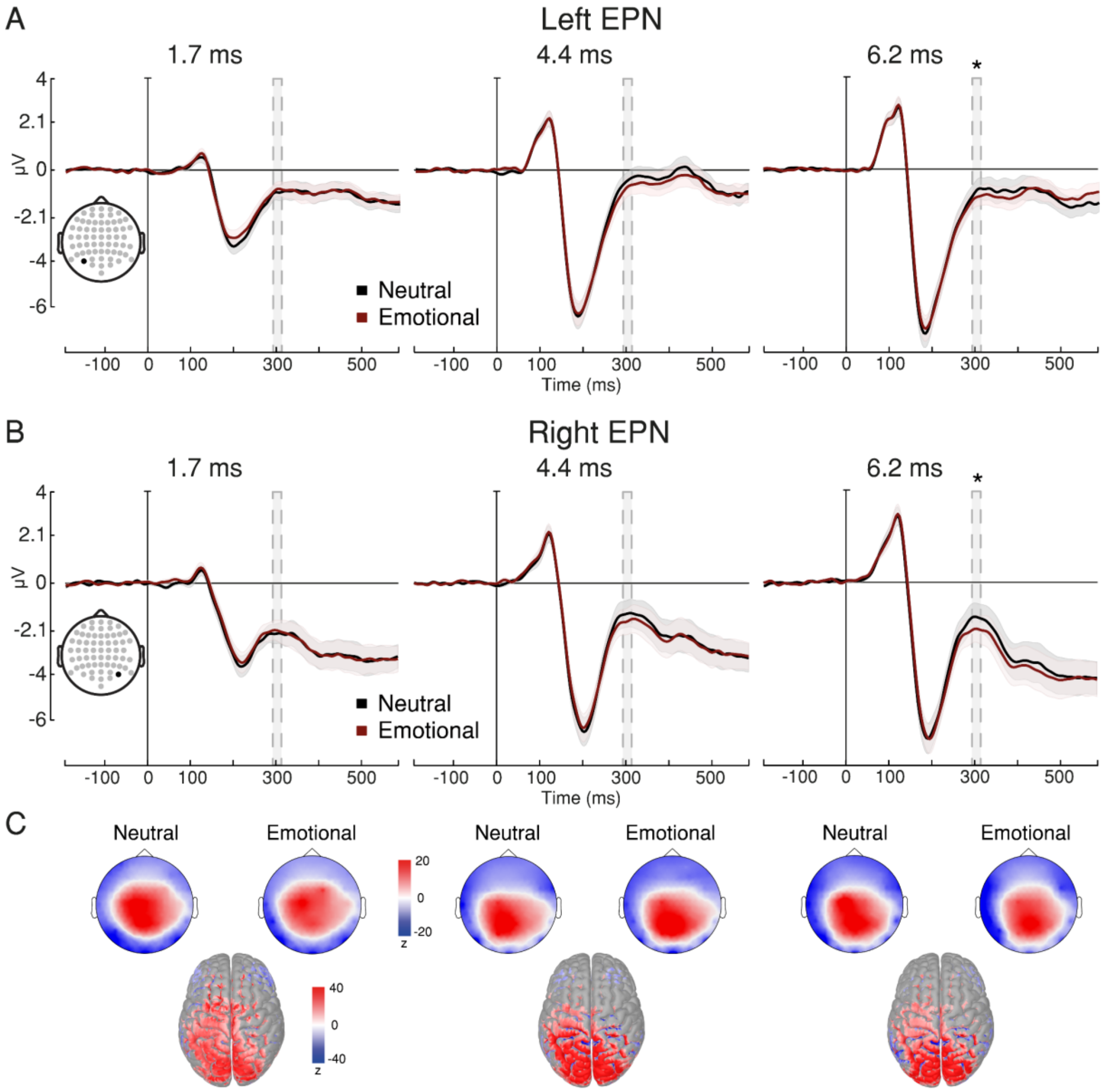


**Supplementary Fig. 20.** Early emotion processing indexed by EPN. (A) Left EPN and (B) right EPN response to emotional and neutral expressions across exposure durations. EPN peak with averaged time window highlighted in grey (295–325 ms). A repeated-measures ANOVA showed that emotional expressions had significantly more negative voltage than neutral expressions at 6.2 ms of exposure. (C) Topographical distributions of EPN for each condition at the relevant time window (z-scale: from -20 to 20). Source estimation of EPN visually identified on cortical maps (z-scale: from -40 to 40). Asterisks indicate statistically significant differences between conditions. Shaded contours represent ±1 SEM, n = 32 independent participants. * $p < 0.05$ for emotional-neutral comparisons.

As an exploratory analysis, we tested whether EPN could discriminate between fearful and happy expressions. To test this, we entered mean voltage values into a 2 (expression: fearful, happy) × 3 (exposure durations) repeated-measures ANOVA. We found a main effect of expression ($F_{(1, 31)} = 8.809, p = .006, \eta p^2 = .221$), indicating that happy expressions had significantly more negative voltage values than fearful expressions. We did not find a main effect of exposure duration ($F_{(1.38, 42.63)} = 2.839, p = .087, \eta p^2 = .084$). The interaction between expression and exposure duration did not reach significance either ($F_{(2, 62)} = 1.905, p = .157, \eta p^2 = .058$). These results might suggest that happy expressions are more emotionally intense given that they are less ambiguous and thus easier to recognise than negative expressions[75–78].

**Late Positive Potential (LPP)**

The late positive potential (LPP) is another ERP component that is sensitive to emotional content and intensity[63,79]. This component is measured over postcentral areas. We entered mean voltage values into a 2 (expression: emotional, neutral) × 3 (exposure durations) repeated-measures ANOVA. We found an effect of exposure duration ($F_{(1.45, 44.97)} = 60.735, p < .00001, \eta p^2 = .662$), indicating that LPP became more positive in voltage as exposure duration increased (**Fig. 3D-F**). We did not find a main effect of expression ($F_{(1, 31)} = 2.119, p = .156, \eta p^2 = .064$) – overall, LPP did not vary between emotional and neutral facial expressions. Crucially, however, we found a significant interaction between expression and exposure duration ($F_{(2, 62)} = 9.804, p = .00019, \eta p^2 = .240$). To test whether LPP was sensitive to emotional content at specific exposure durations, we ran post hoc Bonferroni-corrected pairwise comparisons and found that LPP was significantly more positive for emotional expressions ($M_{6.2\text{ms}}$ = 6.328 [2.878]) than neutral expressions ($M_{6.2\text{ms}}$ = 5.981 [2.904]) only at the longest exposure duration ($t(31) = 4.284$, $p = .00069, d = 0.142, \text{CI} = 0.103 - 0.592$). Thus, these findings of LPP, a marker of late emotion processing, converge with those for the EPN, a marker of early emotion processing, arising by the longest exposure duration used.

As an exploratory analysis, we tested whether LPP could discriminate between fearful and happy expressions. To test this, we entered mean voltage values into a 2 (expression: fearful, happy) × 3 (exposure durations) repeated-measures ANOVA. We found a main effect of exposure duration ($F_{(1.49, 46.1)} = 67.051, p < .00001, \eta p^2 = .684$), suggesting that voltage turned more positive with increasing exposure duration. We did not find a main effect of expression ($F_{(1, 31)} = 0.0533, p = .8188, \eta p^2 = .002$) and the interaction between expression and exposure duration did not reach significance either ($F_{(2, 62)} = 0.074, p = .929, \eta p^2 = .002$). These results might suggest that LPP is sensitive to emotional information relative to neutral information, regardless of emotional valence.

**Visual Awareness Negativity (VAN)**

The visual awareness negativity (VAN) is a voltage difference measured over occipitotemporal areas, bilaterally[70,71,80]. The standard experimental manipulation to extract VAN consists of comparing trials where participants reported seeing a stimulus to those where they reported not seeing it. We examined how awareness, facial expression, and exposure duration conditions affected VAN. By using the distinction between trials with and without awareness based on PAS ratings (see EEG analysis section, above), we could find the exposure-duration range in which visual awareness arises. Since awareness (by including every rating except 'no experience') was present over 90% of trials at the longest exposure duration, we only included trials with the exposure durations of 1.7 ms and 4.4 ms in this analysis (**Supplementary Fig. 21**).

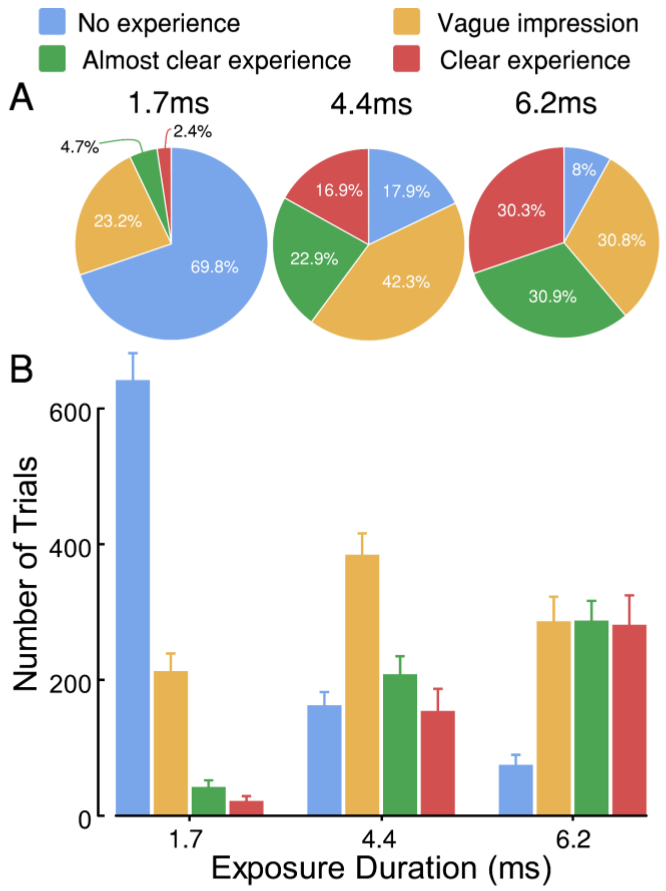


**Supplementary Fig. 21.** Proportion of PAS ratings. (A) Percentages represent overall numbers per exposure duration. At 1.7 ms, 69.8% of the trials were reported as awareness-absent trials ("no experience") and 30.2% were reported as awareness-present trials ("vague impression", "almost clear experience" or "clear experience"). At 4.4 ms, 17.9% of the trials were reported as awareness-absent and 82.1% were reported as awareness-present. At 6.2 ms, 8% of the trials were reported as awareness-absent and 92% were reported as awareness-present. (B) Mean number of trials per PAS rating. Error bars denote ±1 SEM.

We entered mean voltage values into a 2 (awareness: aware, unaware) × 2 (expression: emotional, neutral) × 2 (electrode: left, right) × 2 (exposure durations) repeated-measures ANOVA. We found a main effect of exposure duration ($F_{(1,30)} = 32.042, p = .0000036, \eta p^2 = .516$), indicating that VAN turned more negative with increasing exposure duration (**Supplementary Fig. 22**). Importantly, we found a main effect of awareness ($F_{(1,30)} = 17.755, p = .00021, \eta p^2 = .372$), thus suggesting VAN could discriminate between awareness-present ($M = -4.299$ [1.099]) and awareness-absent trials ($M = -3.535$ [0.825]). Unexpectedly, we also found a main effect of expression ($F_{(1,30)} = 4.427, p = .044, \eta p^2 = .129$), indicating that VAN was significantly less negative in voltage for emotional ($M = -3.747$ [1.073]) than neutral expressions ($M = -4.087$ [1.002]). We did not find an effect of electrode site ($F_{(1,30)} = 1.719, p = .20, \eta p^2 = .054$), indicating that VAN did not significantly vary between hemispheres. Crucially, the interaction between awareness and exposure duration reached significance ($F_{(1,30)} = 10.062, p = .003, \eta p^2 = .251$). To test whether awareness significantly modulated VAN at specific exposure durations, and thus answer our main question here – i.e., at which exposure duration we find a neural indication of awareness (or conscious access) – we ran post hoc Bonferroni-corrected pairwise comparisons and found that VAN was significantly more negative in awareness-present than in awareness-absent trials only at 4.4 ms of exposure ($t(30) = 5.205, p = .000027, d = 0.327, \mathrm{CI} = -1.632 - -0.501$). This finding suggests that 4.4 ms of exposure may be sufficient for faces to reach awareness. We also found a main effect of expression and exposure duration ($F_{(1,30)} = 5.896, p = .021, \eta p^2 = .164$). We did not find significant interactions between expression and awareness ($F_{(1,30)} = 0.194, p = .663, \eta p^2 = .006$), expression and electrode site ($F_{(1,30)} = 4.017, p = .054, \eta p^2 = .118$), electrode site and exposure duration ($F_{(1,30)} = 1.232, p = .276, \eta p^2 = .039$), or any of the three-way interactions and four-way interaction.

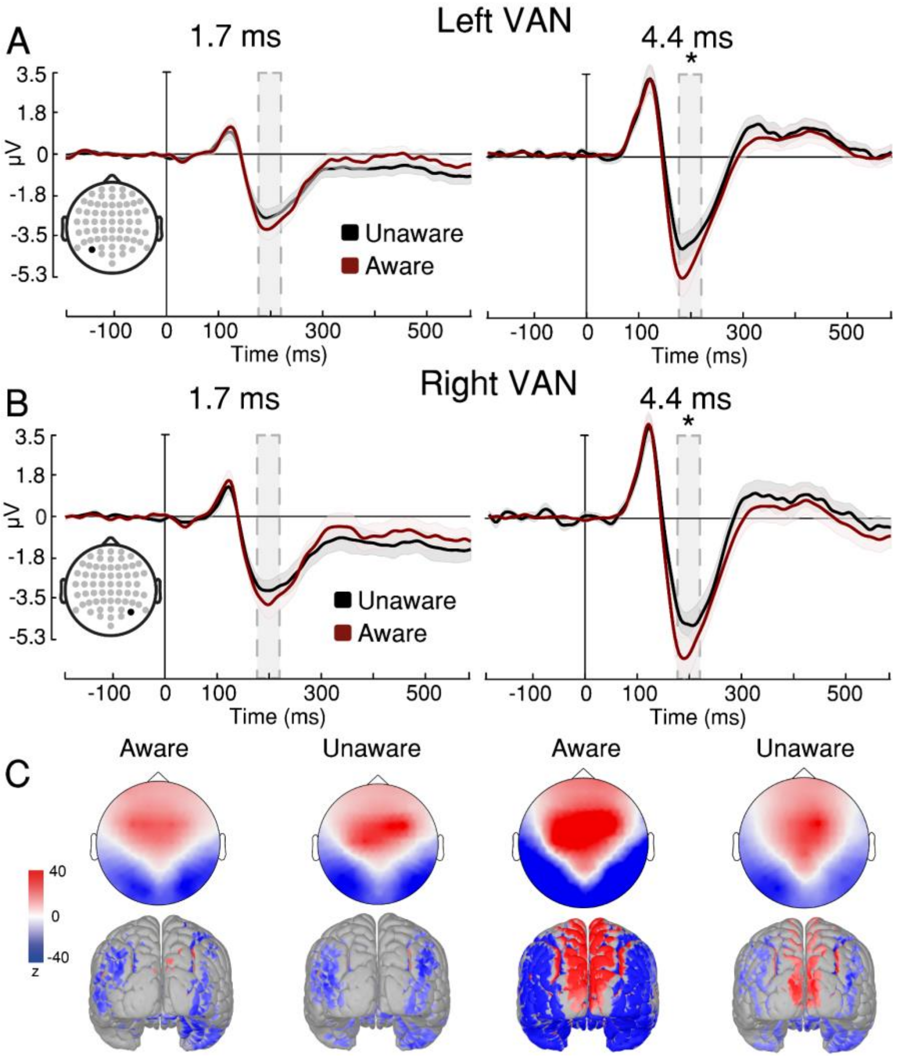


**Supplementary Fig. 22.** Visual awareness indexed by VAN. (A) Left and (B) right VAN response across awareness ratings and exposure durations. Emotional expressions are collapsed together. Averaged time window highlighted in grey (185-215 ms). A repeated-measures ANOVA showed that awareness-present trials had significantly more negative voltage than awareness-absent trials at 4.4 ms of exposure. (C) Topographic maps of VAN (z-scale: from -40 to 40). Source estimations of the ERP at its voltage peak are visually identified on cortical maps (z-scale: from -40 to 40). Shaded contours represent ±1 SEM, n = 32 independent participants. * $p < 0.05$ for aware-unaware comparisons.

### Late Positivity (LP)

The late positivity (LP) is a positive voltage enhancement in the P3 wave. This component is measured over parietooccipital areas, centrally[81–83]. To test whether LP was sensitive to awareness, we sorted trials in the same manner we did with VAN. Then, we entered mean voltage values into a 2 (awareness: aware, unaware) × 2 (expression: emotional, neutral) × 2 (exposure durations) repeated-measures ANOVA. We found a main effect of exposure duration ($F_{(1,30)} = 27.518, p = .000012, \eta p^2 = .478$), indicating that LP became more positive with increasing exposure duration (**Fig. 5C-D**). Importantly, we found a main effect of awareness ($F_{(1,30)} = 7.737, p = .009, \eta p^2 = .205$), with more positive LP voltage in awareness-present ($M$ = 3.597 [0.872]) than awareness-absent trials ($M$ = 3.150 [0.111]). Unexpectedly, we also found a main effect of expression ($F_{(1,30)} = 4.603, p = .04, \eta p^2 = .133$), indicating more positive voltage values for neutral ($M$ = 3.452 [5.590]) than emotional expressions ($M$ = 3.295 [0.736]). This effect is consistent with the (equally unexpected) effect of expression found for the VAN (i.e., greater enhancement for neutral expressions than for emotional ones). Crucially, the interaction between awareness and exposure duration reached significance ($F_{(1,30)} = 37.420, p < .00001, \eta p^2 = .555$). To test at which specific exposure durations LP can discriminate between awareness-present and awareness-absent trials, and thus answer our main question here – i.e., at which exposure duration we find a neural indication of awareness – we ran post hoc Bonferroni-corrected pairwise comparisons and found that LP was significantly more positive in awareness-present than in awareness-absent trials only at 4.4 ms of exposure ($t(55) = 5.861, p = .0000016, d = 0.555, \mathrm{CI} = 0.622 - 1.713$). This finding suggests that 4.4 ms of exposure may be sufficient for enabling conscious access to faces. Importantly, this finding converges with VAN findings. Neither the interaction between expression and awareness ($F_{(1,30)} = 0.457, p = .504, \eta p^2 = .015$) nor the interaction between expression and exposure duration ($F_{(1,30)} = 0.377, p = .544, \eta p^2 = .012$) reached significance. The three-way interaction did reach significance ($F_{(1,30)} = 6.361, p = .017, \eta p^2 = .175$). This interaction seems to be driven by the fact that exposure duration modulated the effect of awareness more strongly for emotional than for neutral expressions. However, we did not examine this in greater detail as we did not have a specific hypothesis concerning such modulations.

#### Discussion of Experiment 5

Are neural systems of emotion processing engaged before faces reach a level of processing that enables expression identification and perceptual awareness? In this experiment, we measured neural markers of visual processing (P1), face processing (N170/VPP), emotion processing (EPN and LPP), and awareness (VAN and LP) with EEG, using three exposure durations that proved relevant according to signal detection analyses in Experiment 1: 1.7 ms (when holistic face processing and expression identification are absent), 4.4 ms (when holistic face processing unfolds), and 6.2 ms (when both expression identification and conscious awareness are very likely to have arisen). Signal detection analyses in Experiment 5 revealed a very similar sequence of processing steps: at 1.7 ms we found weak, but above-chance, stimulus discrimination; and at 4.4 ms above-chance emotion identification, and metacognitive sensitivity arose. By 6.2 ms, all these processes exhibited high sensitivity. If neural markers of emotion processing could distinguish between emotional and neutral expressions at shorter durations than those at which we find behavioural indications of emotion identification, this would provide evidence for unconscious emotion processing. Crucially, we found that both neural markers of emotion processing (EPN and LPP) discriminated between emotional and neutral expressions at 6.2 ms of exposure, meaning that emotion processing arose at some point between 4.4 ms and 6.2 ms of exposure. Together, neural markers and signal detection indices suggest that emotion processing does not arise before holistic face processing and awareness do.

How much visual exposure is required for faces to reach perceptual awareness? To address this question, we measured two neural markers of awareness (VAN and LP); both markers discriminated between awareness-present and awareness-absent trials when faces were presented for 4.4 ms, thus suggesting that perceptual awareness requires between 1.7 ms and 4.4 ms of visual exposure, alongside holistic face processing (as indexed by the FIE of location d′ in Experiment 1) and expression identification (as shown in Experiment 1 and 2). However, what aspects of awareness VAN and LP specifically index is still a matter of debate[71,84]. Some studies have suggested that VAN may index phenomenal consciousness (i.e., subjective experience) whereas LP would index conscious access (i.e., the ability to consciously access and thus report sensory information[70,72,82,85–87]). Regardless of this distinction's validity, the fact that both neural markers discriminated between awareness-present and awareness-absent trials by 4.4 ms of exposure may suggest that perceptual awareness emerges alongside face holistic processing. It is interesting to note that we also found an unexpected effect of emotion on both VAN and LP, which might suggest that these components are affected by other cognitive processes, too. For example, researchers using Binocular Rivalry, found that VAN was modulated by awareness and emotional content, which they interpreted as evidence of an emotional bias in perceptual awareness[88]. It has been also suggested that VAN, N170, and EPN may be affected by different but correlated underlying processes when studying the interaction between face perception, emotion processing, and awareness, as those three markers share very similar topographies and time windows[89]. Future studies should expand on whether VAN specifically indexes awareness or other aspects involved in perception that may be necessary but not sufficient for awareness. Similarly, the effect of emotion on LP might have been driven by the LPP's sensitivity to emotion content; LP and LPP have similar topographies and time windows, and both are part of the P3 wave[51,90].

Crucially, however, two multiclass Multivariate pattern analyses (MVPA; one of emotional expression, and one of intact-face location) found evidence of emotion processing by 4.4 ms of exposure, thus suggesting that emotion processing may have begun unfolding as holistic face processing did (indexed by the FIE of location d′ in Experiment 1). As shown in past studies[91], MVPA is sensitive to multivariate patterns that ERP analysis is not; this may explain why MVPA here found emotion processing with a shorter exposure duration than EPN and LPP did.

So far, however, all experiments have used only images of faces. To what extent are the processes and durations we found specific to faces? In Experiment 1, we found that holistic processing, indexed by the FIE in location sensitivity, required a minimal exposure duration of 4.4 ms to unfold. Do neural systems of face processing require the same visual exposure to discriminate a face from a non-face stimulus? In Experiment 5, we measured the N170/VPP complex, which is a neural marker of face processing. However, we did not include a control condition to determine by which exposure duration neural face processing arises. To determine the minimal required exposure of face processing, we need to measure the N170/VPP in response to face and non-face stimuli.

**Supplementary Note 12: Full results of Experiment 6**

In Experiment 5, we used signal detection indices and EEG neural markers to test whether emotion processing arises before perceptual awareness. While with ERP analysis we found that neural emotion processing may have arisen with 6.2 ms of exposure, with multiclass MVPA we found that it may have arisen with 4.4 ms of exposure (**Fig. 4**), thus suggesting that emotion processing begins unfolding as faces gain access to awareness. Therefore, perceptual awareness may be required for emotion processing to occur. But is this the case of face processing? Does face processing arise before perceptual awareness does? It could be the case that while emotion processing may require perceptual awareness to arise, face processing arises before perceptual awareness does.

Here, our main question is whether neural systems can discriminate face from non-face stimuli before holistic processing and perceptual awareness arise. In Experiment 1, we found evidence of holistic face processing – indexed by the FIE in location sensitivity – by 4.4 ms of exposure for stimuli presented peripherally. But may neural markers that are specific to face processing be evident even before holistic processing (a hallmark of specialised face processing) arises? In Experiment 5, we measured the N170 component, which is sensitive to face and face-like visual information. The N170 component belongs to the N1-family, a group of visually evoked potentials that are elicited over visual cortical areas in response to visual stimulation[43,48,92]. N170 (and the whole N170/VPP ERP complex) exhibits a significantly more negative voltage evoked to face stimuli than non-face stimuli. Since in Experiment 5 we did not have non-face stimuli to employ as controls, we could not tell whether the responses of N170 in were specific to faces as any kind of visual processing could explain it. By comparing the ERP evoked by face and non-face stimuli we can find the exposure duration at which the response evoked by faces becomes a face sensitive N170, distinguishable from the N1 evoked by non-face stimuli. If N170 can discriminate between faces and non-faces before there is sufficient visual information available to elicit the FIE (i.e., around 4.4 ms), then neural systems can process facial information before holistic face processing unfolds. On the other hand, if the same minimal exposure duration is required to generate an N170 that discriminates between face and non-face stimuli as is required for the FIE to arise, then face processing may indeed require holistic processing to occur.

**Behavioural results**

**Location sensitivity**

We measured location sensitivity to confirm that our new participants displayed comparable performance to participants in the equivalent conditions of Experiment 1. To examine how conditions affected stimulus discrimination, we entered location d′ scores into a 2 (stimulus category: face, object) × 4 (exposure durations) repeated-measures ANOVA. We found a main effect of exposure duration ($F_{(1.28, 39.75)} = 184.675, p < .00001$, ηp² = .856), indicating that location d′ scores increased with increasing exposure duration (**Supplementary Fig. 23A**). We also found a main effect of stimulus category ($F_{(1, 31)} = 5.629, p = .024$, ηp² = .154), with higher location d′ scores for objects ($M = 0.709$ [0.777]) than faces ($M = 0.567$ [0.751]). Finally, the interaction between the two factors also reached significance ($F_{(2.06, 63.8)} = 8.921, p < .00034$, ηp² = .223). To test whether location d′ scores differed between stimulus categories at any exposure duration, we ran post hoc Bonferroni-corrected pairwise comparisons and found significantly higher location d′ scores for objects over faces at 2.45 ms of exposure ($t(31) = 5.386, p < .000014, d = 1.059$, $\mathrm{CI} = -0.752 - -0.190$). This advantage, however, may be due to low-level visual differences such as stimulus size – no object could fill the oval entirely like faces, which may have led to contrast differences. Overall, these results indicate that location sensitivity to both faces and objects increases with increasing exposure duration even in the brief range from 0.8 to 4.288 ms. Importantly, the location d′ scores we found resemble the ones found in Experiment 1.

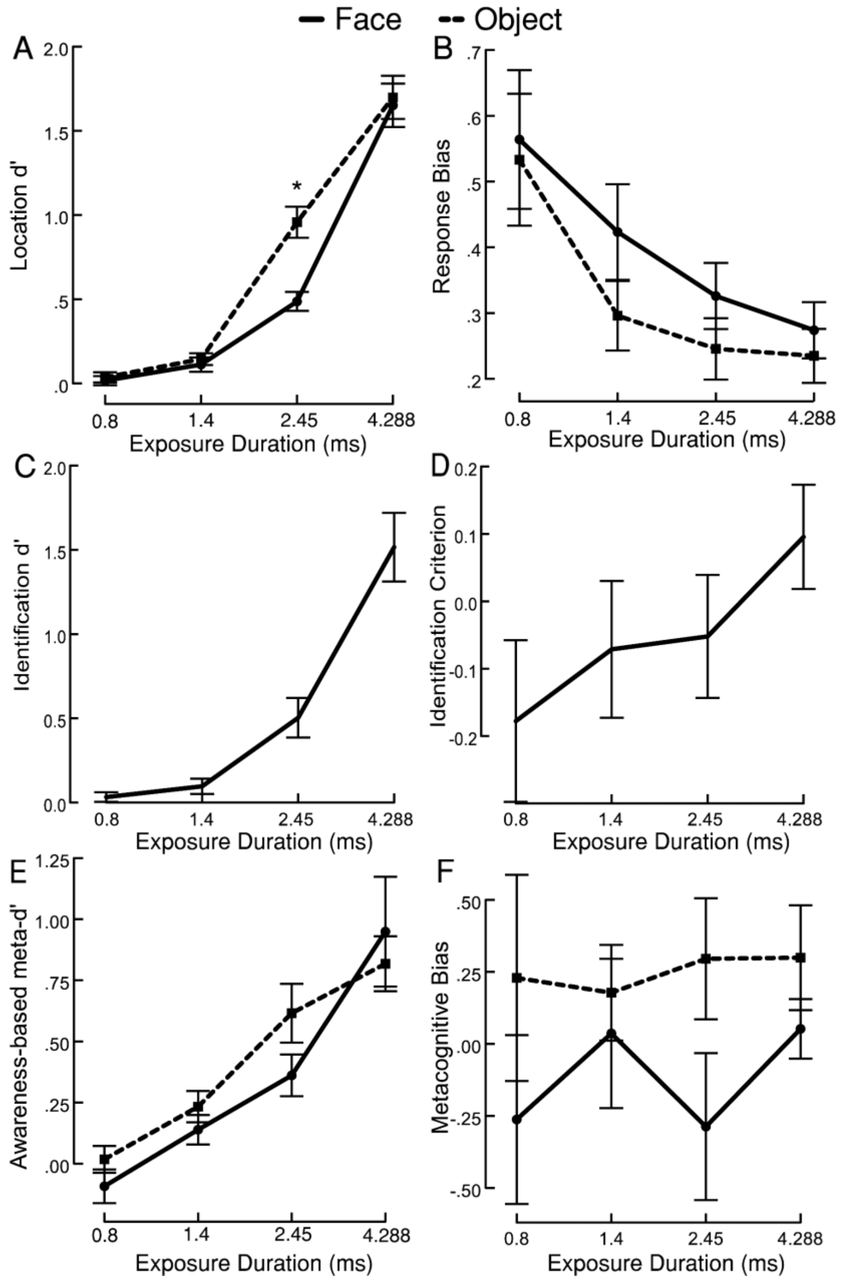


**Supplementary Fig. 23.** Full behavioural results of Experiment 6. (A) Location sensitivity. A repeated-measures ANOVA showed that location d′ increased with increasing exposure duration. A significant advantage for objects over faces was found at 2.45 ms of exposure. (B) Absolute-value location response bias scores for reporting location (bias toward either left or right). A repeated-measures ANOVA showed that the amount of bias decreased as exposure duration increased, with greater bias for faces than objects. (C) Identification sensitivity for stimulus category. A repeated-measures ANOVA showed that participants' ability to discriminate faces and objects increased with exposure duration. (D) Criterion scores for reporting stimulus category. A repeated-measures ANOVA showed that lower criterion indicates greater bias to reporting a face. Criterion does not significantly change with increasing exposure duration. (E) Awareness-based metacognitive sensitivity. A repeated-measures ANOVA showed that meta-d′ increased with exposure duration but was unaffected by stimulus category. (F) Metacognitive bias scores for reporting subjective awareness. A repeated-measures ANOVA showed that metacognitive bias was unaffected by exposure duration and stimulus category. Data are presented as mean values with ±1 SEM bars, n = 32 independent participants. * $p < 0.05$ for face-object comparisons.

To determine the minimal required exposure that exhibited above-chance performance (d′ > 0), we ran a series of uncorrected one-sample t-tests against zero. We found that the earliest exposure duration that elicited above-chance discrimination was 1.4 ms for both face ($M = 0.112$ [0.239]; $t(31) = 2.65, p = .013, d = 0.468$. CI $= 0.026 - 0.198$) and object stimuli ($M = 0.145$ [0.201]; $t(31) = 4.08, p = .00029, d = 0.72$, CI $= 0.072 - 0.217$).

#### Location response bias

We examined whether participants' response bias for reporting stimulus location varied across conditions by entering the absolute values of $C_{identification}$ scores into a 2 (stimulus categories: face, object) × 4 (exposure durations) repeated-measures ANOVA. As in previous experiments, response bias significantly decreased with exposure duration (**Supplementary Fig. 23B**), as indicated by a main effect of exposure duration ($F_{(1.276, 39.56)} = 8.255, p = .004, \eta p^2 = .210$). We also found a main effect of stimulus category ($F_{(1, 31)} = 8.59, p = .006, \eta p^2 = .217$), indicating higher amount of response bias for faces ($M = 0.387$ [0.127]) than objects ($M = 0.328$ [0.140]). The interaction between these two factors did not reach significance ($F_{(3, 93)} = 1.794, p = .154, \eta p^2 = .055$).

#### Stimulus category identification sensitivity

We examined whether participants' sensitivity to discriminating faces from objects varied across exposure durations by comparing identification d′ scores in a one-way ANOVA (**Supplementary Fig. 23C**). As expected, identification d′ increased alongside exposure duration ($F_{(3, 54.794)} = 32.428, p < .00001, \eta p^2 = .44$).

To determine the minimal required exposure that exhibited above-chance performance (d′ > 0), we ran a series of uncorrected one-sample t-tests against zero. We found that the earliest exposure duration that elicited above-chance stimulus category identification was 1.4 ms ($M = 0.096$ [0.260]; $t(31) = 2.09, p = .045, d = 0.369$, CI $= 0.002 - 0.190$).

#### Stimulus category identification criterion

We examined whether participants' criterion for reporting faces rather than objects varied across exposure durations by comparing criterion scores in a one-way ANOVA (**Supplementary Fig. 23D**). Lower scores indicated a greater tendency to report a face. Identification criterion did not vary across exposure durations ($F_{(3, 54.791)} = 32.429, p < .00001, \eta p^2 = .44$).

#### Awareness-based metacognitive sensitivity

We examined whether awareness scores were sensitive to participants' location sensitivity scores by calculating meta-d′. To examine whether metacognitive sensitivity varied across conditions, we entered meta-d′ scores into a 2 (stimulus category: face, object) × 4 (exposure durations) repeated-measures ANOVA (**Supplementary Fig. 23E**). A main effect of exposure duration indicated that meta-d′ increased with increasing exposure duration ($F_{(1.79, 55.36)} = 24.20, p < .00001, \eta p^2 = .438$). However, we did not find a main effect of stimulus category

($F_{(1,31)} = 0.902, p = .350, \eta p^2 = .028$), suggesting that stimulus category did not affect metacognitive sensitivity. We did not find an interaction between stimulus category and exposure duration either ($F_{(2.06, 63.96)} = 1.744, p = .182, \eta p^2 = .053$).

As described, we did not find a main effect of stimulus category, therefore we calculated Bayes factors to test whether the obtained data support this absence of an effect. Bayes factors indicated substantial evidence in favour of the null hypothesis model ($BF_{01} = 4.67$), supporting the finding that meta-d′ was not greater for either stimulus category.

To determine the minimal exposure that exhibited above-chance performance (meta-d′ > 0), we ran a series of uncorrected one-sample t-tests against zero. We found that the earliest exposure duration that elicited above-chance metacognitive awareness was 1.4 ms for both face ($M = 0.139$ [0.341]; $t(31) = 2.304, p = .028, d = 0.407, \mathrm{CI} = 0.016 - 0.262$) and object stimuli ($M = 0.233$ [0.362]; $t(31) = 3.643, p = .00098, d = 0.644, \mathrm{CI} = 0.103 - 0.364$).

**Metacognitive bias**

As described above, meta-bias is the tendency to give high confidence (or awareness) ratings regardless of actual performance. To examine whether meta-bias varied across conditions, we entered meta-bias scores into a 2 (stimulus category: face, object) × 4 (exposure durations) repeated-measures ANOVA (**Supplementary Fig. 23F**). We did not find an effect of exposure duration ($F_{(2.396, 74.288)} = 0.679, p = .536, \eta p^2 = .021$), or of stimulus category ($F_{(1,31)} = 0.76, p = .39, \eta p^2 = .024$). The interaction between stimulus category and exposure duration did not reach significance either ($F_{(2.158, 66.895)} = 1.459, p = .239, \eta p^2 = .045$). These results suggest that the participants' tendency to describe their subjective awareness was not affected by any of the conditions.

**EEG results**

**P1**

We examined how stimulus categories and exposure duration conditions affected early visual processing by measuring voltage changes in P1. This component is measured over occipital regions. We entered mean voltage values into a 2 (stimulus category: face, object) × 4 (exposure durations) repeated-measures ANOVA. We found a main effect of exposure duration ($F_{(1.52, 47.02)} = 81.863, p < .00001, \eta p^2 = .725$), whereby P1 amplitude increased with increasing exposure duration, indicating that longer exposure durations involve more visual processing than shorter exposure durations (**Supplementary Fig. 24**). We did not find a main effect of stimulus category, suggesting that images of faces and objects did not affect early visual processing differently ($F_{(1,31)} = 1.489, p = .232, \eta p^2 = .046$). The interaction between these factors reached significance ($F_{(3,93)} = 2.885, p = .04, \eta p^2 = .085$), but post hoc Bonferroni-corrected pairwise comparisons did not reveal significant differences between stimulus categories at any exposure duration. These results indicate that P1 was sensitive to visual differences between extremely brief exposure durations.

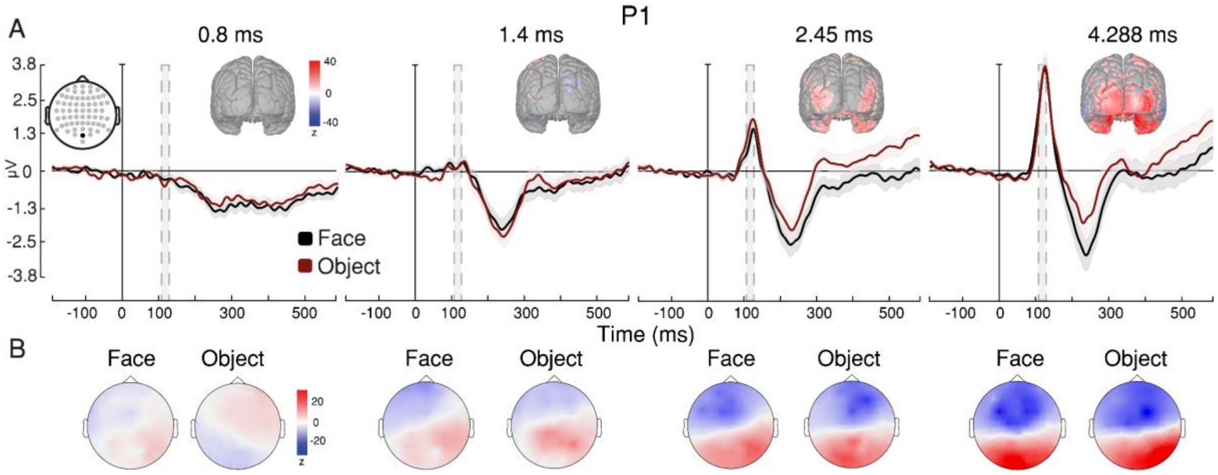


**Supplementary Fig. 24.** Early visual processing indexed by P1. (A) P1 evoked response to face stimuli is represented by black lines, while evoked response to object stimuli is represented by red lines, across exposure durations. P1 peak with averaged time window highlighted in grey (105–135 ms) and source estimation of P1 visually identified on cortical maps (z-scale: from -40 to 40). (B) Topographical distributions of P1 for each condition at the relevant time window show an increase in positive voltage over visual cortex across exposure durations (z-scale: from -50 to 50). Shaded contours represent ±1 SEM, n = 32 independent participants.

### N170/VPP

We examined how stimulus category and exposure duration conditions affected face processing by measuring voltage changes in N170/VPP[43,92,93]. This component is measured over occipitotemporal regions, bilaterally, and over frontocentral regions. We entered mean voltage values into a 2 (stimulus categories: face, object) × 3 (electrode site: left, right, central) × 4 (exposure durations) repeated-measures ANOVA. We found a main effect of exposure duration ($F_{(1.39,\,42.92)} = 45.681, p < .00001, \eta p^2 = .596$), indicating that the magnitude of this component significantly increased with increasing exposure duration. We also found a main effect of electrode site ($F_{(1.31,\,40.594)} = 62.566, p < .00001, \eta p^2 = .669$), which was expected given that left ($M = -3.225$ [1.510]) and right N170 peaks ($M = -3.394$ [1.603]) are negative in voltage whereas VPP is positive ($M = 2.789$ [1.050]). Like in Experiment 5, this effect helped confirm that we were measuring the N170/VPP complex. We did not find a main effect of stimulus category ($F_{(1,\,31)} = 0.005, p = .946, \eta p^2 < 0.01$), but crucially, we found a significant interaction between stimulus category and exposure duration ($F_{(2.45,\,75.89)} = 6.398, p = .001, \eta p^2 = .171$). To test whether N170/VPP was sensitive to faces at specific exposure durations, we ran post hoc Bonferroni-corrected pairwise comparisons and found that the N170/VPP was significantly greater in magnitude for face stimuli ($M_{\mathrm{N170:\,4.288ms}} = -5.093$ [0.307]) compared to object stimuli ($M_{\mathrm{N170:\,4.288ms}} = -4.290$ [0.159]) only at the longest exposure duration ($t(31) = 3.467, p = .021, d = 0.134, \mathrm{CI} = -0.668 - -0.027$), thus indicating that 4.288 ms of visual exposure conveyed sufficient visual information for neural systems to distinguish between face and non-face stimuli (**Fig. 6** and **Supplementary Fig. 25**). These results suggest that neural systems require around the same visual exposure to trigger face processing as is required for the FIE to arise (FIE in location d'; Experiment 1, main text).

Taken together, these findings indicate that neural face processing requires 4.288 ms of exposure – and definitely no less than 2.45 ms where we see no difference.

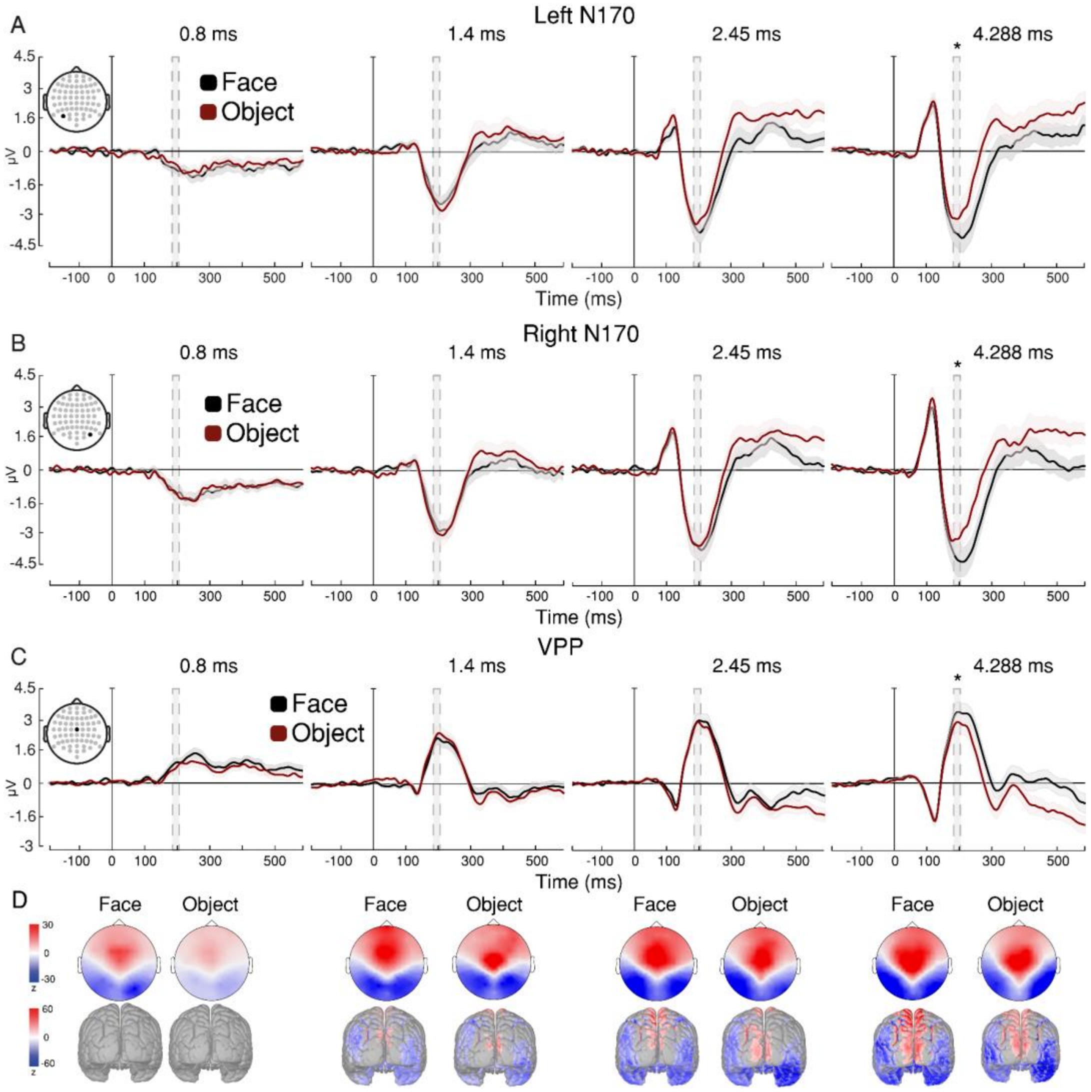


**Supplementary Fig. 25.** Face processing indexed by N170/VPP. (A) Left N170, (B) right N170, and (C) VPP response across exposure durations. N170/VPP evoked response to face stimuli is represented by black lines, while evoked response to object stimuli is represented by red lines. N170/VPP peak with averaged time window highlighted in grey (175–205 ms). A repeated-measures ANOVA showed that the magnitude of N170/VPP increased with exposure durations, and Bonferroni-corrected post hoc pairwise comparisons showed an advantage for faces with 4.288 ms of exposure. (D) Topographical distributions of N170/VPP for each condition at the relevant time window (z-scale: from -30 to 30) and source estimations visually identified on cortical maps (z-scale: from -60 to 60). Shaded contours represent ±1 SEM, n = 32 independent participants. * $p < 0.05$ for face-object comparisons.

While the N170/VPP complex has a specific topographic distribution, latency, and neuroanatomical source[50,94], **Fig. 6** seems to suggest different evoked voltage responses to face and object images at later latencies, with shorter exposure durations. We conducted an exploratory analysis of the voltage responses for all four exposure durations at later latencies, by taking the mean voltage values between 300 and 500 ms after stimulus onset. To test whether the small numerical difference between means (see **Fig. 6**) is statistically significant, we entered these new mean voltage values into a 2 (stimulus categories: face, object) × 3 (electrode site: left, right, central) × 4 (exposure durations) repeated-measures ANOVA. We found a main effect of exposure duration ($F_{(1.36,\,42.17)} = 12.827, p = .00027, \eta p^2 = .293$), indicating a significant change in mean voltage as exposure duration increased. We also found a main effect of stimulus category ($F_{(1,\,31)} = 14.956, p < .00053, \eta p^2 = .325$), which was expected. Most importantly, we found a significant interaction between stimulus category and exposure duration ($F_{(2.25,\,69.78)} = 3.06, p = .047, \eta p^2 = .09$). However, none of the relevant post-hoc pairwise comparisons between faces and objects revealed significant differences: at 0.8 ($t(31) = 0.171, p = 1, d = 0.005, \mathrm{CI} = -0.231 - 0.209$), 1.4 ($t(31) = 1.864, p = 1, d = 0.079, \mathrm{CI} = -0.526 - 0.155$), 2.45 ($t(31) = 3.367, p = .057, d = 0.161, \mathrm{CI} = -0.766 - 0.006$), or 4.288 ms of exposure ($t(31) = 2.844, p = .219, d = 0.122, \mathrm{CI} = -0.632 - 0.058$), (**Supplementary Fig. 26**). We also found that the interactions between stimulus category and electrode ($F_{(1.52,\,47.09)} = 22.998, p < .00001, \eta p^2 = .426$), and the three-way interaction between stimulus category, electrode, and exposure duration ($F_{(4.5,\,139.43)} = 4.291, p = .002, \eta p^2 = .122$) were significant. However, no post-hoc pairwise comparison of interest reached significance.

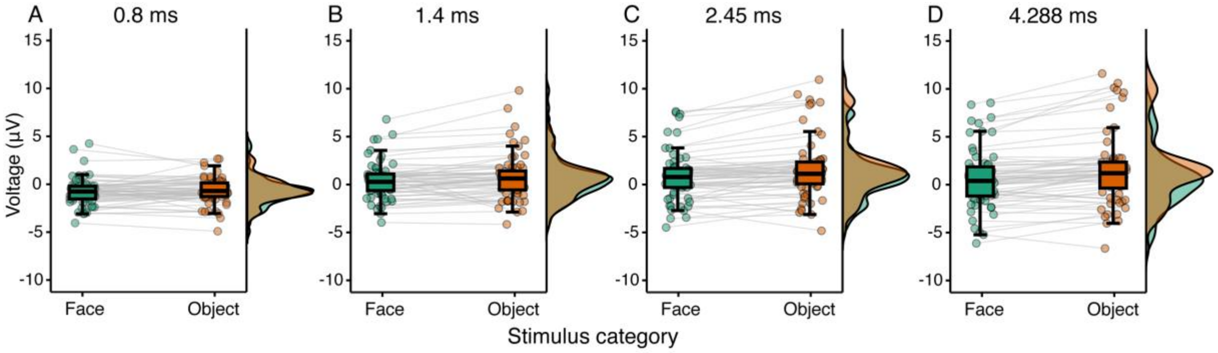


**Supplementary Fig. 26.** Exploratory analysis of late latencies in the visual evoked potential across exposure durations. Mean voltage values between 300 and 500 ms after stimulus onset recorded at P07 and PO8 electrode sites (i.e., the same electrodes used for capturing the left and right N170 components, respectively). Evoked response to face stimuli is represented in green, and for object stimuli is represented in orange. A repeated-measures ANOVA found no significant differences were found with (A) 0.8, (B) 1.4, (C) 2.45, or (D) 4.288 ms of exposure. Boxplots show median and interquartile range with individual dot plots and their distribution.

### Visual Awareness Negativity (VAN)

We examined whether VAN could distinguish between awareness-present and awareness-absent trials across exposure durations by sorting trials the same way we did in Experiment 5. This component is measured over occipitotemporal regions, bilaterally. We entered mean voltage values into a 2 (awareness: aware, unaware) × 2 (stimulus category: face, object) × 2 (electrode: left, right) × 4 (exposure durations) repeated-measures ANOVA. We found a main effect of exposure duration ($F_{(1.94,\,58.11)} = 18.86, p < .0001, \eta p^2 = .386$), indicating that VAN became more negative with increasing exposure duration (**Supplementary Fig. 27**). We also found a main effect of stimulus category ($F_{(1,\,30)} = 4.258, p = .048, \eta p^2 = .124$), which indicated that faces evoked more negative voltage means ($M = -2.607$ [1.076]) than objects ($M = -2.270$ [0.931]). This effect may be due to the fact that VAN was measured at the same electrode sites as N170, and at a later yet close time window. Importantly, we found a main effect of awareness ($F_{(1, 30)} = 19.612, p < .00012, \eta p^2 = .395$) – awareness-present trials had more negative voltage means ($M = -2.766$ [1.044]) than awareness-absent trials ($M = -2.112$ [0.880]). Importantly, the interaction between awareness and exposure duration did not reach significance in this experiment ($F_{(3, 90)} = 1.368, p = .258, \eta p^2 = .044$), suggesting that VAN distinguished between awareness-present and awareness-absent trials overall, but in a linear fashion. This finding is in line with the view that VAN may be an index of phenomenal consciousness – arguably, such a marker should be sensitive to awareness reports regardless of conscious access to sensory information. Therefore, if VAN were a marker of phenomenal consciousness, it should correlate with awareness reports only in a linear way. Interestingly, the interaction between stimulus category and exposure duration reached significance ($F_{(2.49, 74.81)} = 3.708, p = .021, \eta p^2 = .110$). To test whether VAN was sensitive to stimulus categories at specific exposure durations, we ran post hoc Bonferroni-corrected pairwise comparisons and found that VAN was significantly more negative for faces than objects only at the longest exposure duration: 4.288 ms ($t(30) = 3.877, p = .005, d = 0.375, \mathrm{CI} = -2.066 - -0.199$). This effect may have been driven by voltage changes in N170, as this component and VAN share electrode sites and have very similar temporal windows. We did not find a main effect of electrode site ($F_{(1, 30)} = 1.845, p = .185, \eta p^2 = .058$), suggesting that VAN changes did not differ between hemispheres. Finally, we did not find significant interactions between awareness and electrode site ($F_{(1, 30)} = 1.096, p = .303, \eta p^2 = .035$), stimulus category and awareness ($F_{(1, 30)} = 0.930, p = .343, \eta p^2 = .03$), electrode site and exposure duration ($F_{(1.95, 58.41)} = 1.269, p = .288, \eta p^2 = .041$), nor any three-way interaction and the four-way interaction. In conclusion, VAN was sensitive to awareness ratings provided by participants, irrespective of visual exposure.

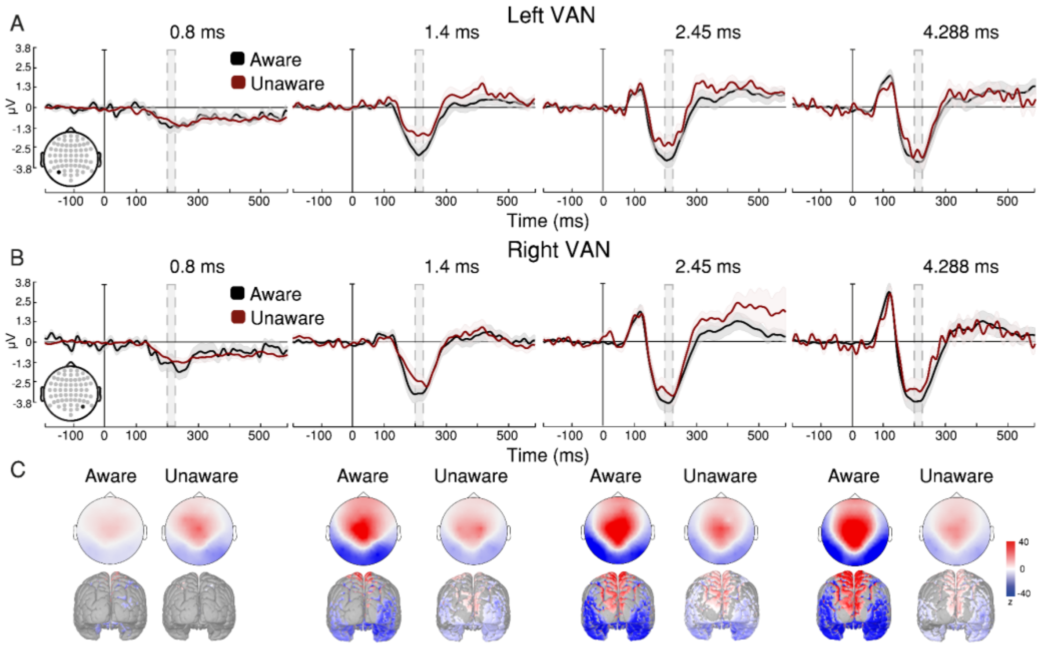


**Supplementary Fig. 27.** Visual awareness indexed by VAN. (A) Left and (B) right VAN response across awareness ratings and exposure durations. Stimulus categories are collapsed together. Evoked responses associated with aware reports are represented by black lines, while evoked responses associated with unaware reports are represented by red lines. Averaged time window highlighted in grey (200-230 ms). A repeated-measures ANOVA found that evoked responses significantly decreased across exposure durations, but no interaction between exposure duration and awareness report was found. (C) Topographic maps of VAN (z-scale: from -4 to 4). Source estimations of the ERP at its voltage peak are visually identified on cortical maps (z-scale: from -40 to 40). Shaded contours represent ±1 SEM, n = 31 independent participants.

### Late Positivity (LP)

We examined whether LP could distinguish between awareness-present and awareness-absent trials by sorting trials the same way we did in Experiment 5. This component is measured over parietooccipital regions. We entered mean voltage values into a 2 (awareness: aware, unaware) × 2 (stimulus category: face, object) × 4 (exposure durations) repeated-measures ANOVA. We found a main effect of exposure duration ($F_{(1.99,\,59.78)} = 11.715, p = .000052, \eta p^2 = .281$), indicating that LP voltage increased with increasing exposure duration (**Supplementary Fig. 28**). Importantly, we found a main effect of awareness ($F_{(1,\,30)} = 4.826, p = .036, \eta p^2 = .139$), which indicates that LP voltage was significantly more positive in awareness-present trials ($M = 2.621\ [0.858]$) than in awareness-absent trials ($M = 2.142\ [0.372]$). Therefore, like in Experiment 5, LP was sensitive to awareness ratings. Crucially, the interaction between awareness and exposure duration reached significance, indicating that exposure duration may have significantly modulated LP's sensitivity to awareness ($F_{(2.34,\,70.034)} = 5.393, p = .004, \eta p^2 = .152$). To test whether LP was sensitive to awareness ratings at specific exposure durations, we ran post hoc Bonferroni-corrected pairwise comparisons and found that LP was significantly more positive in awareness-present than awareness-absent trials only at the longest exposure duration: 4.288 ms ($t(30) = 3.683, p = .011, d = 0.529, \mathrm{CI} = 0.140 - 2.097$). This finding suggests that neural systems require around said visual exposure to unfold conscious access. We did not find a main effect of stimulus category ($F_{(1,\,30)} = 0.168, p = .685, \eta p^2 = .006$) nor significant interactions between stimulus category and awareness ($F_{(1,\,30)} = 2.069, p = .161, \eta p^2 = .065$), and between stimulus category and exposure duration ($F_{(3,\,90)} = 0.319, p = .811, \eta p^2 = .011$). The three-way interaction between awareness, stimulus category, and exposure duration was not significant either ($F_{(2.32,\,69.66)} = 1.764, p = .174, \eta p^2 = .056$).

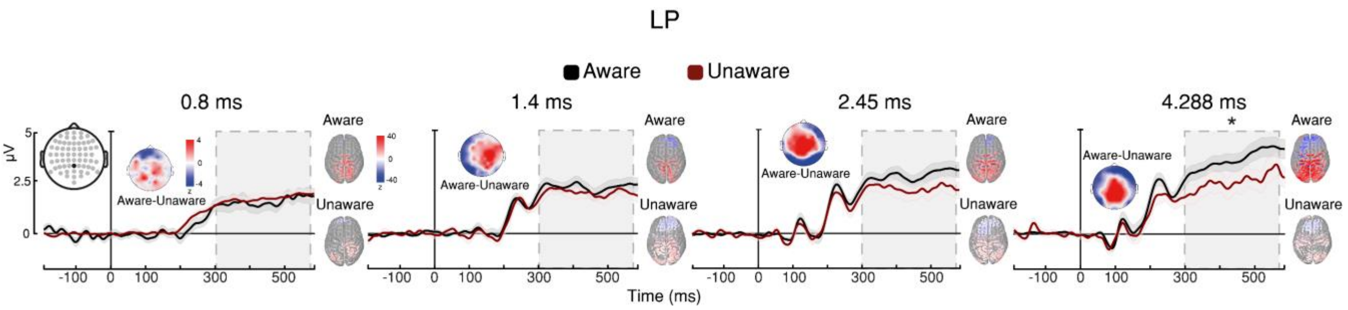


**Supplementary Fig. 28.** Visual awareness indexed by LP. LP response across awareness ratings and exposure durations. Stimulus categories are collapsed together. Evoked responses associated with aware reports are represented by black lines, while evoked responses associated with unaware reports are represented by red lines. Averaged time window highlighted in grey (300-585 ms). A repeated-measures ANOVA found that evoked response significantly increased with exposure duration, and Bonferroni-corrected post hoc comparisons revealed a higher evoked response in aware-report trials compared to unaware-report trials. Topographic maps of LP (z-scale: from -4 to 4). Source estimations of the ERP at its voltage peak are visually identified on cortical maps (z-scale: from -40 to 40). Shaded contours represent ±1 SEM, n = 31 independent participants. * $p < 0.05$ for aware-unaware comparisons.

**Discussion of Experiment 6**

In Experiment 6, we measured neural markers of face processing to test whether they can discriminate face and non-face stimuli at shorter exposures than signal detection indices do. We found that neural markers could discriminate between faces and objects at 4.288 ms of exposure, but not at shorter durations. This exposure duration roughly corresponds to the minimal exposure duration required for the FIE, as shown in Experiment 1. Therefore, our findings using neural markers here converge with findings from signal detection indices in Experiment 1, indicating that around 4.4 ms of visual exposure are needed for neural systems to engage in face-specific processing.

What is the minimal exposure duration required for faces to access perceptual awareness? We found convergent evidence of faces (and objects) gaining access to awareness by 4.288 ms of exposure coming from signal detection indices (metacognitive sensitivity) and neural markers of awareness and conscious access (VAN and LP). Importantly, unlike in Experiment 5 where both VAN and LP discriminated between awareness-present and awareness-absent trials by 4.4 ms, in Experiment 6 only LP interacted with exposure duration and exhibited discrimination between awareness-present and awareness-absent trials specifically by 4.288 ms of exposure. Although VAN exhibited a main effect of awareness, it did not interact with exposure duration. Importantly, these findings may suggest that faces gain access to awareness as their visual information becomes available for holistic processing.

Some studies have suggested that VAN indexes phenomenal consciousness (i.e., changes in subjective experience) whereas LP indexes conscious access[71,72,80,82]. Our findings in Experiment 6 may support this distinction and provide evidence of a minimal amount of visual exposure required for faces to gain access to awareness. Evidence from the N170/VPP complex and from signal detection analyses in this experiment converge, thereby suggesting that faces require around 4.288 ms of exposure to gain access to perceptual awareness; this duration is similar to that required for holistic face processing in Experiment 1. This temporal convergence may suggest an underlying causal relationship between holistic face processing and awareness – e.g., awareness may be necessary for integration of visual features to allow or facilitate face recognition and subsequent emotional processing.

In summary, we found evidence with neural markers that face processing begins unfolding by around 4.288 ms of exposure. Similarly, we found evidence both with neural markers and signal detection indices that perceptual awareness of faces unfolds by around 4.288 ms of exposure as well. Together, these findings suggest that face processing and conscious access may unfold together.

**Supplementary Note 13: The relationship between stimulus energy and information**

Our primary research question focused on the processing priorities of the visual system and their relation to awareness. Some claims suggest that specific types of visual information, such as the configurations that make a face, or that make an expression emotional, are prioritised by the visual system. Previous attempts to assess this claim, however, have been hindered by the need to use visual masks. Our core methodological advancement is the introduction of a way to present stimuli so briefly, using the tachistoscope, that they do not require masks to be below, on, and just above the thresholds for perception and awareness, thus eliminating the confounding effects of masking.

The key results derived from this advancement are twofold: (1) a sequence of required durations that unveils the priorities in the visual system, and (2) a demonstration that this sequence is influenced by one manipulation (face inversion) but not by another (emotional expression).

Using displays of varying exposure durations (a form of bottom-up stimulation) is a manipulation of different levels of information accumulation. Therefore, the minimal exposure durations we identified provide upper bounds specific to the stimuli used. We would expect stimuli with higher luminance or greater distinctiveness to require shorter minimal exposures while still exhibiting the same order of required durations we found.

Nonetheless, the role of stimulus energy at very brief exposure durations is not a settled question. Bloch's famous finding that the "*visibility of a candlelight is markedly in inverse proportion to its duration*"[95], has historically been interpreted as a rule of temporal summation – "*if the brightness of a stimulus is halved, the stimulus can still be detected if its duration is doubled*"[96]. However, the validity of Bloch's law for brief presentations of the type used here has been called into question. For instance, recent studies by Greene[97,98] have suggested that Bloch's law may not hold for stimulus displays in the range of 10 µs to 10 ms, particularly in tasks involving shape recognition. Moreover, investigations into temporal dynamics in feature vision have also provided evidence contradicting Bloch's law, showing that energy summation alone cannot explain feature vision[99]. Finally, the research supporting Bloch's law has been conducted with very simple stimuli, such as spots[100,101] and thin bars[102], rather than complex images containing multiple high-level features. As such, it is unclear how the law generalises to more complex stimuli, especially concerning the extraction of meaning rather than mere detection of the presence of something. For instance, we discovered different minimal exposures for upright and inverted faces, despite their identical energy; clearly, Bloch's law cannot account for this.

To delve deeper into the relationship between stimulus energy and meaning extraction, particularly for complex stimuli like faces, further studies will need to combine manipulations of both extremely brief display durations and stimulus energy. An important goal should be to understand the degree to which the processing priorities of the visual system can be explained in terms of the information or energy present in a stimulus, versus the particular goals of human vision.
